# *NET1* mRNA localization to the midbody is required for Arp2/3-dependent initiation of mitotic abscission

**DOI:** 10.64898/2026.08.07.743554

**Authors:** Katherine F. Vaeth, Andrew J Neumann, Frances Zorensky, Xinrui Wei, Rytis Prekeris, J. Matthew Taliaferro

## Abstract

The recruitment and activation of abscission machinery following mitosis is tightly spatiotemporally regulated, yet the mechanisms underlying this remain poorly understood. We find that RNA localization and local translation at the midbody governs when and where abscission-regulating proteins are expressed. The 3′ UTR of *NET1* mRNA contains an element that is necessary and sufficient for RNA targeting to the midbody. Mislocalization of *NET1* mRNA results in a loss of NET1 protein, a Rho family GEF, throughout the intercellular bridge, delaying abscission and slowing cell proliferation. Arp2/3-mediated branched actin accumulation at the midbody, which is necessary for abscission, requires localized NET1 protein that is competent for binding Rho family GTPases. These findings establish midbody RNA localization and local translation as a key regulatory layer over abscission timing and identify a role for NET1 as a regulator of Arp2/3-mediated branched actin accumulation at the abscission site.

## INTRODUCTION

Abscission is the last step of mitotic cell division that leads to physical separation of two daughter cells. This highly regulated process requires the sequential recruitment and activation of specific proteins at defined subcellular domains (Elia et al. 2011; Mierzwa and Gerlich 2014; Neto and Gould 2011). Among the key abscission regulators are Rho family GTPases that govern the spatiotemporal actin dynamics necessary for cleavage furrow establishment, ingression, and abscission (Haga and Ridley 2016; David et al. 2012; Jaffe and Hall 2005; Schmidt et al. 2007). The ingression of the cleavage furrow drives the formation of the intercellular bridge (ICB) connecting the dividing cells (Basant and Glotzer 2018; Wagner and Glotzer 2016). At the center of the ICB, the central spindle microtubules are crosslinked in an anti-parallel fashion to form a midbody (MB) (D’Avino and Capalbo 2016; Hu et al. 2012).

The MB is critical in regulating the timing and location of abscission, although the molecular machinery governing this process remains to be fully understood. In addition to tightly packed microtubules and abscission regulating proteins, the MB contains a specific subset of localized mRNAs (Farmer et al. 2023; Park et al. 2023) that encode several known abscission-regulating proteins, such as CHMP4B and Citron kinase (Farmer et al. 2023; Park et al. 2023). Importantly, MB-enriched mRNAs are specifically localized to the MB and are distinct from both mitotically enriched or spindle enriched mRNAs, suggesting the existence of specific molecular machinery that mediates targeting of these distinct mRNAs to the MB. However, it remains unknown how MB-enriched mRNAs are localized and whether their localization to the MB is important for regulation of abscission (Farmer et al. 2023).

Thousands of RNAs are known to be asymmetrically distributed within cells. The localized translation of these mRNAs often facilitates the spatiotemporal regulation of protein localization and function and is essential for diverse processes such as cell migration, neuronal polarization, and development (Costa et al. 2020; Holt and Bullock 2009; Macdonald and Struhl 1988; Ryder and Lerit 2018; Gasparski et al. 2023; Lawrence and Singer 1986). However, the molecular machinery governing mRNA localization to specific subcellular domains remains to be fully defined.

The subcellular distribution of mRNAs is often regulated by localization elements (LEs) within the 3’UTR of the mRNA itself. These elements are recognized by trans-factors to facilitate RNA trafficking to specific subcellular compartments (Arora et al. 2022; Wang et al. 2023; Macdonald and Struhl 1988). Although thousands of RNAs display distinct subcellular localization patterns, the localization elements and trans-factors responsible for this distribution are known for only a small number of them (Engel et al. 2020).

Furthermore, known examples of cellular defects arising from the mislocalization of specific mRNAs remain rare, particularly in mammalian cells. This knowledge gap exists, at least in part, because perturbing the localization of a specific RNA often requires knowledge of the mechanisms (e.g. localization elements and trans-factors) that control its transport.

In this study, we identify NET1 mRNA, which encodes a Rac/Rho guanine nucleotide exchange factor (GEF), as localized to and translated at the MB during abscission. We characterize the LE within NET1 mRNA that mediates its targeting to the MB and demonstrate that disruption of this LE inhibits abscission by preventing the accumulation of NET1 protein at the ICB during late telophase. Finally, we show that the loss of MB-localized NET1 mRNA, and therefore MB-localized NET1 protein, disrupts Arp2/3-dependent branched actin accumulation at the abscission site, likely through the loss of Rac1 signaling. Together, these findings establish NET1 mRNA localization to the MB as important for abscission and identify NET1 protein as a new regulator of Arp2/3-dependent branched actin formation at the abscission site, a step that promotes faithful completion of mitosis. Our work also identifies mRNA targeting and local translation at the ICB as a key process that contributes to the regulation of timing and location of the abscission site during mitotic division.

## RESULTS

### A sequence element within the 3’UTR of NET1 mRNA is necessary and sufficient for its localization to the midbody

The midbody contains a specific population of enriched mRNAs, but it remains poorly understood how these mRNAs get to the MB or whether their localization is important for cell cycle progression and abscission (Farmer et al. 2023; Park et al. 2023). To understand the role of MB localized mRNAs, we focused on *NET1*, an mRNA that is strongly and reproducibly localized both to the MB and to other subcellular locations in a variety of cell types (Farmer et al. 2023; Suwakulsiri et al. 2024; Goering et al. 2023; Gasparski et al. 2023; Park et al. 2023). Although LEs within *NET1* mRNA that are responsible for its transport to cellular protrusions and the basal pole of epithelial cells have been identified, how *NET1* mRNA is trafficked to MB remains unknown (Engel et al. 2020; Arora et al. 2022; Mason et al. 2025; Gasparski et al. 2023). Further, it is not known what role, if any, NET1 protein plays during abscission.

RNA localization elements are often found in 3’UTRs (Martin and Ephrussi 2009; Engel et al. 2020; Arora et al. 2022). We therefore first tested whether *NET1* mRNA localization to the MB is dependent on its 3’UTR. We leveraged the fact that membrane-bound MBsomes (also referred to as MB remnants or Flemmingsomes) are released into the extracellular space from cells that undergo symmetric abscission (Peterman et al. 2019; Crowell et al. 2014; Skop et al. 2004; Patel et al. 2024). These released MBsomes can then be purified using a series of ultracentrifugation steps based on their specific size and density (Peterman and Prekeris 2017).

Localization to the MB can then be tested using a system in which two reporter RNAs are transcribed from a bidirectional promoter (**Fig S1A**). We site-specifically integrated this system into HeLa cells and used qRT- PCR to quantify the relative abundance of the two reporter RNAs in MB and whole cell (WC) RNA samples. By comparing the ratio of the two reporter RNAs in the MB and WC samples, we can quantify their relative enrichment in MBs (**Fig S1A**). We can then append specific sequences to one of the two reporter RNAs to ask how their inclusion in an RNA affects its localization to MBs.

Relative to a control 3′UTR, we found that the inclusion of the 3’UTR of *NET1* was sufficient to drive a reporter RNA to the MB (**Fig 1A**). However, as the full-length 3’UTR sequence is quite large, we next asked what sequences within the 3’UTR are responsible for this localization activity. To address this, we used a massively parallel reporter assay (MPRA) to interrogate the localization capacity of hundreds of sequences drawn from the full length *NET1* 3’UTR.

**Figure 1.**
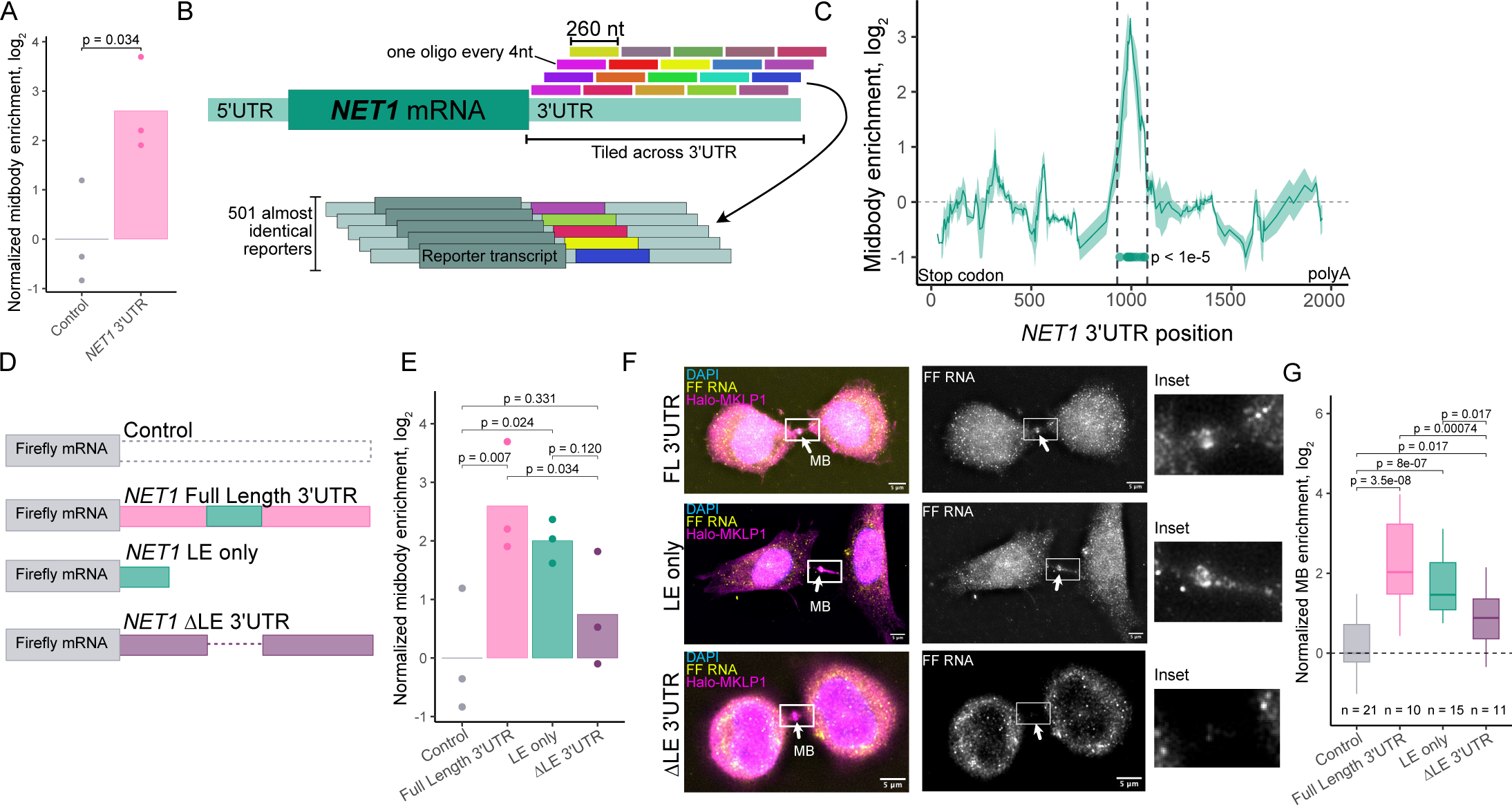
The identification of an RNA localization element within the NET1 3’UTR. (A) Midbody enrichment (log_2_) of a reporter RNA appended to the NET1 full length 3’UTR, normalized to the reporter with a control, plasmid-derived 3’UTR appended. The bar represents the mean midbody enrichment of independent biological replicates (dots). P values were calculated using a t-test. (B) Design of an MPRA to identify RNA elements that mediate RNA transport to the midbody. Oligonucleotides of length 260 nt were designed against the 3′ UTR of MB localized genes. Oligonucleotides were spaced 4 nt from each other, giving an average coverage of 65X per nucleotide. Each oligonucleotide was then integrated into the 3′ UTR of a reporter transcript, generating about 501 nearly identical reporters to assay MB localization. (C) Midbody enrichment of reporter-embedded oligonucleotides as a function of their location within the *NET1* 3’UTR. The line represents a rolling average of 8 oligonucleotides and the ribbon represents the standard deviation of MB enrichment within the sliding window. Dots below the lines represent 3’UTR positions of significantly MB localized oligos (FDR < 1e-5). Dotted lines indicate the location of the *NET1* localization element. (D) Schematic of reporter RNA constructs used to assess the necessity and sufficiency of the identified NET1 localization element. (E) Midbody localization of the reporter RNAs in D quantified using a midbody purification and qPCR (Fig S1. A). Values are normalized to the control reporter. P values were calculated using a t test. (F) smFISH visualizing the reporter RNAs described in D (yellow) in cells stably expressing Halo-MKLP1 (pink). Midbodies are indicated by arrows. The box indicates the area shown in a higher magnification inset. (G) Quantification of NET1 reporter RNAs visualized in F. Reporter transcript counts in intercellular bridges/MBs and whole cells were quantified for each sample and the ratio between the two locations is reported as normalized midbody enrichment (log_2_). Values were normalized to the control reporter. P values were calculated using a t test.

We designed 260-nucleotide oligonucleotides (oligos) tiled across the entire 3’UTR of human *NET1* mRNA, with neighboring oligos shifting by four nucleotides. This resulted in approximately 500 partially overlapping oligos spanning the entire length of the human *NET1* 3’UTR (**Fig 1B, S1B**). Given that we had previously identified an element in the mouse *Net1* RNA sufficient for RNA transport to MBs (Farmer et al. 2023), we also included oligos that tiled the mouse *Net1* 3’UTR as a positive control.

Each oligo was then cloned into the 3′UTR of the exact same reporter construct, thus creating a library of reporter RNAs whose only difference between them lay in the identity of the integrated oligo. Therefore, differences in MB localization between the reporters could be attributed to the specific oligo it contained (**Fig 1B**). These reporters were site-specifically integrated into HeLa cells using a *cre*/loxP system such that every cell received only one reporter RNA from the library.

We then isolated MB and WC RNA samples as before and used targeted high-throughput sequencing to quantify the relative abundances of oligos in the MB and whole cell RNA samples (Arora et al. 2022).

Encouragingly, a PCA analysis of oligo abundances revealed a clear separation between WC and MB samples, indicating a reproducible difference in their composition (**Fig S1C**). We found that oligos drawn from a discrete region within the *NET1* 3’UTR were significantly more enriched in the MB RNA compared to whole cell RNA, indicating that they were sufficient to drive the reporter RNA to the MB (**Fig 1C**). We defined this region of the 3’UTR as the *NET1* localization element (LE).

To further characterize this new, putative LE, we next asked whether the *NET1* LE is both necessary and sufficient to drive MB localization. Using our previously described qRT-PCR-based reporter system (**Fig S1A**), we appended either the full-length *NET1* 3’UTR, the full length UTR with the LE deleted (ΔLE), or the LE alone to the reporter (**Fig 1D**). We found that the LE alone was sufficient to drive midbody localization of this reporter RNA at a level similar to the full length UTR (**Fig 1E**). Deleting the LE resulted in a loss of reporter RNA localization to the MB, indicating its necessity for MB localization (**Fig 1E**). These findings were further validated using single molecule fluorescence *in situ* hybridization (smFISH) with fluorescently labeled probes designed against the open reading frame of the reporter transcript. We again observed MB localization of reporter RNAs containing either the full-length 3’UTR or the LE alone to the MB and a loss of MB localization for the ΔLE reporter RNA, further confirming that the *NET1* LE identified using the MPRA is both necessary and sufficient for MB RNA localization (**Fig 1F, G**).

### Antisense oligos designed against the NET1 LE disrupt NET1 mRNA localization to the MB and inhibit cell proliferation

An antisense phosphorodiamidate morpholino oligonucleotide (PMO) developed against a region in the *NET1* 3’UTR has been previously shown to block the localization of *NET1* RNA to the plus-end of microtubules (Gasparski et al. 2023). The *NET1* LE we identified overlaps with the sequence previously targeted by PMOs to mislocalize *NET1* in epithelial cells (Mason et al. 2025; Gasparski et al. 2023). Thus, we reasoned that this PMO may also be an efficient tool to begin to investigate if *NET1* mRNA localization to the MB is required for mitotic cell division.

First, using smFISH against endogenous *NET1*, we asked if treatment with a PMO designed against the LE (LE PMO) would mislocalize *NET1* mRNA away from the MB and ICB. As controls, we designed two other PMOs. One PMO (scramble control) contained a scrambled sequence of the LE PMO. A second PMO (3′UTR control) targeted the *NET1* 3′UTR but at a region outside of the LE (**Fig 2A**). Treatment of HeLa cells with either of the control PMOs did not result in mislocalization of *NET1* mRNA away from the MB. In contrast, *NET1* mRNA was robustly mislocalized in the LE PMO treated cells (**Fig 2B-D**), further suggesting that this LE is involved in targeting *NET1* mRNA to the MB during mitosis.

**Figure 2.**
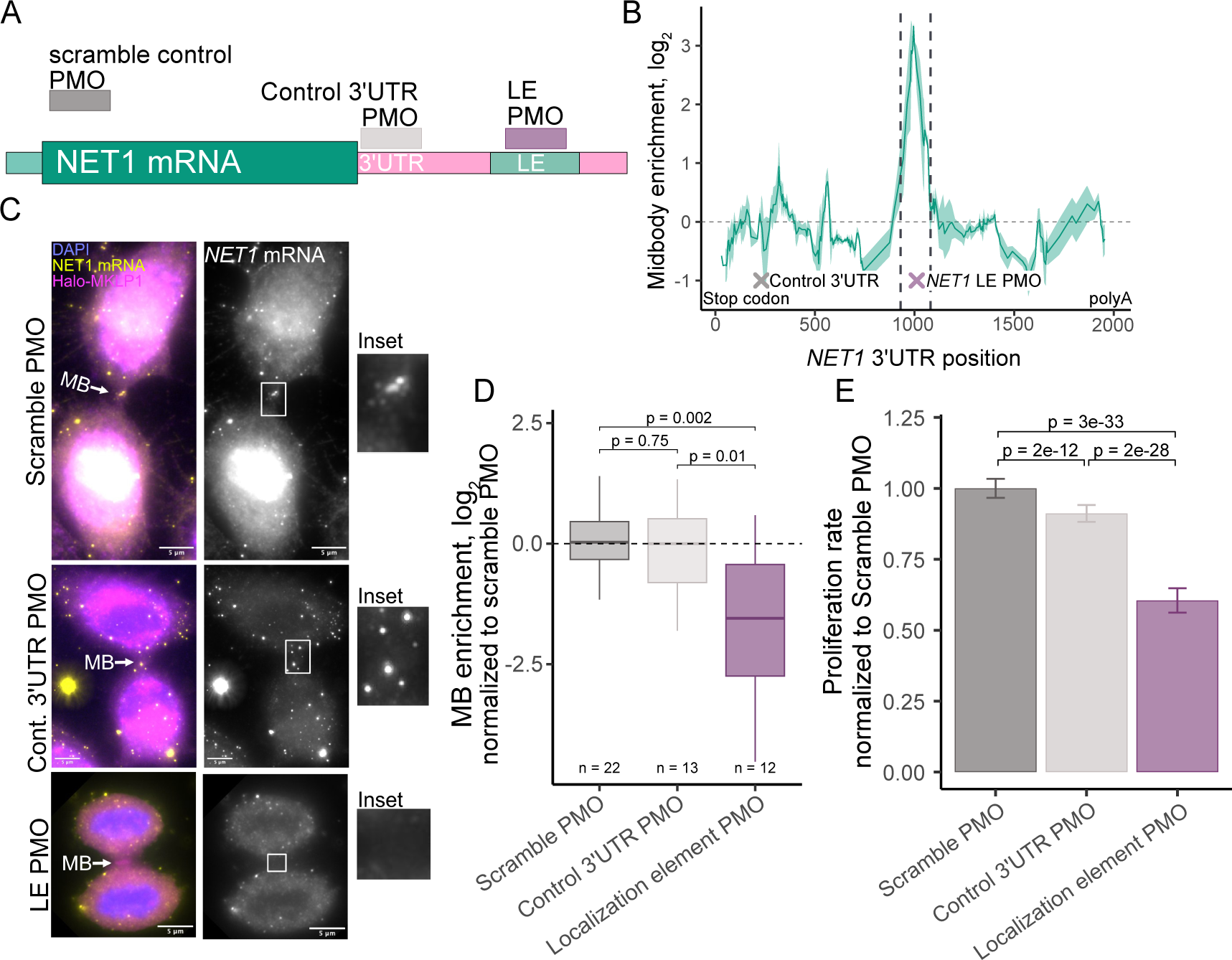
Antisense oligos designed against the NET1 LE disrupt NET1 mRNA localization to the MB and inhibit cell proliferation. (A) Schematic of NET1 showing the regions targeted by phosphorodiamidate morpholino oligonucleotides (PMOs). (B) MB enrichment of oligonucleotides spanning the 3’UTR of *NET1* overlaid with X’s indicating regions targeted by PMOs. (C) Midbody localization of *NET1* (yellow) with different PMO treatments visualized using smFISH in cells stably expressing Halo-MKLP1 (pink). Midbodies are indicated by arrows. The box indicates the area shown in a higher magnification inset. (D) Quantification of *NET1* mRNA visualized in C. *NET1* puncta in ICBs/MBs and whole cells were quantified for each sample and the ratio between the two locations is reported as normalized midbody enrichment (log_2_). Values were normalized to the scramble control. P values were calculated using a t test. (E) Proliferation rate for cells treated with PMOs, normalized to scramble PMO. Percent confluency was measured every 3 hours over a period of 96 hours.

To test whether *NET1* mRNA localization to the MB is important during cell division we next analyzed cell proliferation in the absence or presence of the LE PMO. We observed that cells treated with the LE PMO proliferated significantly slower than cells treated with either of the control PMOs (**Fig 2E**). Importantly, PMO treatment did not affect *NET1* mRNA levels (**Fig S1E**), indicating that the effect of LE PMO treatment on cellular proliferation is likely due to *NET1* mRNA mislocalization rather than a change in the overall *NET1* mRNA expression or stability.

### Localization of NET1 mRNA to the MB is necessary for NET1 protein accumulation at the ICB during late telophase

To further define the role of *NET1* mRNA MB localization during cell division, we turned to a system which would allow us to study a homogenous population of cells in which *NET1* mRNA has been mis- localized rather than the heterogenous PMO-treated cell population. We therefore generated a NET1 knockout HeLa cell line using CRISPR-Cas9 and two guide RNAs that targeted early and late exons in the open reading frame of *NET1* (**Fig S2A**). We isolated single cell clones and screened these clones via PCR, identifying a clone where at least one allele had lost all sequence between the gRNA cut sites and all other alleles contained a frameshift-generating single nucleotide deletion (**Fig S2B-D**).

We then rescued this knockout line using doxycycline-inducible transgenes that were site-specifically integrated into the genome (Khandelia et al. 2011). These transgenes encoded HA-tagged NET1 protein followed by either the full length *NET1* 3′UTR (FL 3′UTR) or a version of this 3′UTR in which the LE has been deleted (ΔLE 3′UTR) (**Fig 3A**). Importantly, we found that both rescue constructs were expressed at similar levels and did so in a doxycycline-dependent manner (**Fig 3B**). Therefore, the phenotypic differences observed between the two rescues are likely due to the localization of the mRNA and not due to differences in protein expression or degradation.

**Figure 3.**
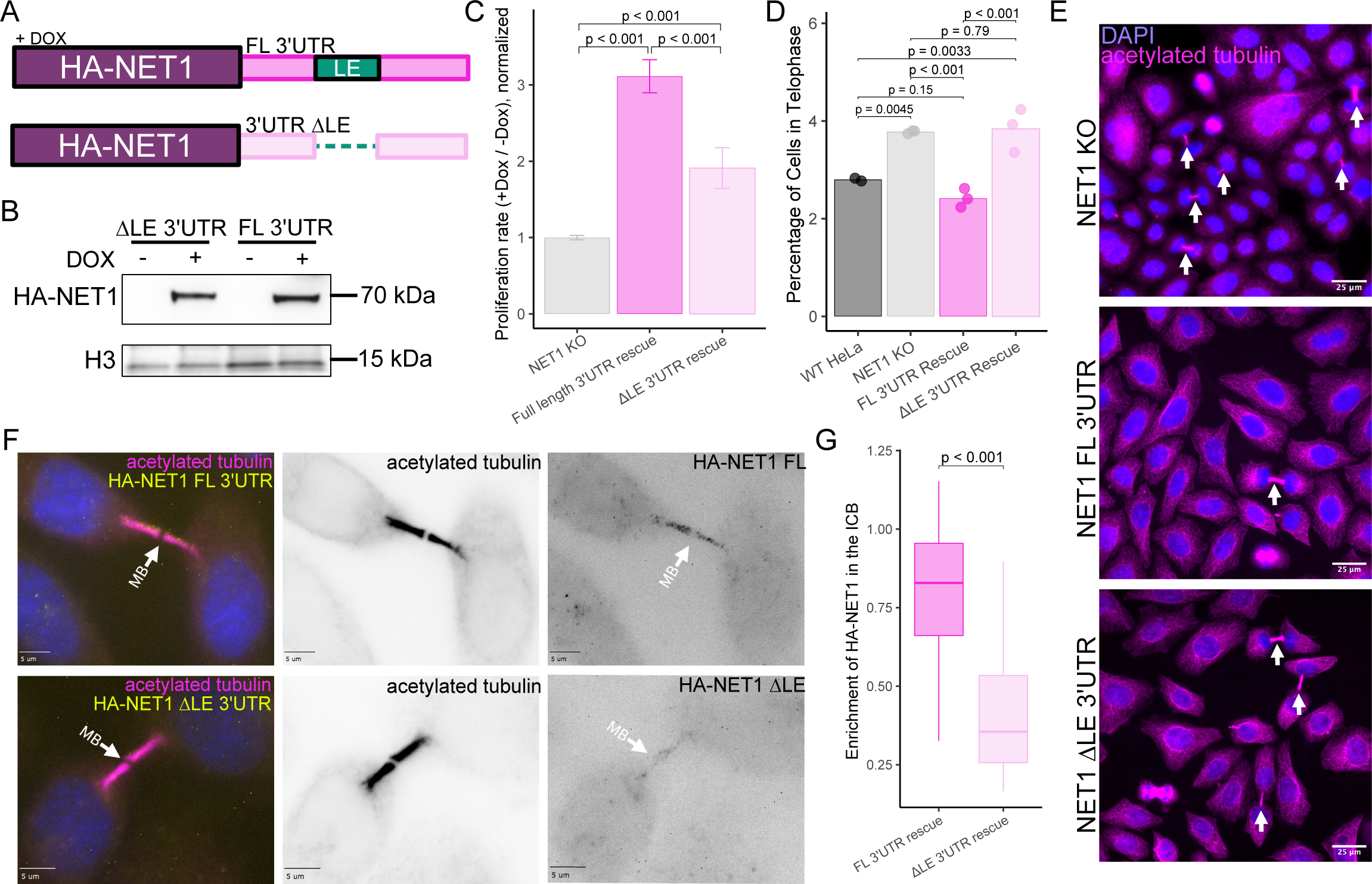
Localization of NET1 mRNA to the MB is necessary for NET1 protein accumulation at the ICB during late telophase. (A) Schematic of doxycycline inducible HA-NET1 rescue constructs used to assess how localization affects proliferation defects. (B) Western blot measuring protein expression in HA-NET1-FL 3’UTR and HA-NET1-ΔLE 3’UTR expressing cell lines with and without doxycycline. Histone 3 (H3) is used as a loading control. (C) Proliferation rate analysis for NET1 KO cells and lines expressing the HA-NET1 rescue constructs. Values observed in plus-doxycycline (transgene expressed) conditions are normalized to those seen in minus-doxycycline (transgene not expressed) conditions. Percent confluency was measured every 3 hours over a period of 96 hours. P values were calculated using a t test. (D) Percentage of cells in telophase identified using DAPI and anti-acetylated tubulin immunofluorescence. Each point represents a biological replicate with a minimum of 1000 cells represented. P values were calculated using a t-test. (E) Representative immunofluorescence images for quantification shown in panel C. Telophase cells were identified by presence of an intercellular bridge stained with anti-acetylated tubulin antibody (magenta). Arrows indicate cells in telophase. (F) HA-NET1 (yellow) localization during late telophase. Cells in late telophase exhibit rounded nuclei (blue), and fully formed ICBs (magenta). Midbodies are indicated by arrows. (G) Quantification of HA-NET1 enrichment in the ICB as represented by the images in F. P values were calculated using a t-test.

We next tested localization of rescue HA-NET1 mRNA during cell division. FL 3′UTR transgene mRNA localized to the MB, which is consistent with our previous data that 3′UTR is sufficient to localize NET1 to the MB (**Fig S2E-G**). In contrast, the ΔLE 3′UTR transgene mRNA was not localized to the MB (**Fig S2E-G**) further demonstrating that *NET1* LE is required for *NET1* mRNA accumulation at the MB.

Our NET1 knockout and dox-inducible rescue cell lines allow us to test whether NET1 is required for mitotic cell division, especially abscission. NET1 has previously been implicated in regulating mitotic spindle assembly but its role during abscission remains unknown (Menon et al. 2013; Wang et al. 2024; Ulu et al. 2021). This system allows us to test whether *NET1* mRNA and NET1 protein localization to the ICB is required for mitotic NET1 function.

First, we analyzed whether NET1 KO affects overall cell proliferation rates. Unfortunately, we observed large variability in proliferation rates among the individual NET1 knockout/rescue lines, likely due to clonal variation. Because of this variability, we were unable to directly compare proliferation rates of individual KO clones to those of a parental HeLa cell line. To control for this clonal variation, we instead normalized proliferation rates within rescue lines between the doxycycline-plus (transgene expression) and doxycycline-minus (no transgene expression) conditions. Consistent with our hypothesis that NET1 is required for cell division, rescue with the FL 3′UTR transgene resulted in a higher proliferation rate as compared to NET1 KO cells (**Fig 3C**). Using the ΔLE 3′UTR transgene only partially rescued proliferation defects suggesting that LE-dependent mRNA targeting to the MB is essential for mitotic NET1 function (**Fig 3C**).

A decrease in proliferation rates can be caused by a variety of defects in the cell cycle. Indeed, it has been previously reported that NET1 may be important during mitotic spindle formation and positioning (Wang et al. 2024; Menon et al. 2013), although the role of NET1 during abscission was never investigated. To further demonstrate that *NET1* mRNA targeting is important during mitosis, we quantified the percentage of cells in metaphase or telophase using an acetylated-tubulin antibody, a known marker for mitotic spindle and ICB microtubules, since an accumulation of cells in a specific mitotic stage can be an indication of the defects in that state of cell division.

In the NET1 KO cells, we observed a distinct increase in the portion of cells in both metaphase and telophase (**Fig 3D,E, S2H**), suggesting that cells may be defective in both mitotic spindle assembly and abscission. We found that expression of the FL 3’UTR transgene rescued the accumulation of cells in both metaphase and telophase (**Fig 3D,E, S2H**). Rescuing with ΔLE 3′UTR transgene, however, did not decrease the number of cells in telophase (**Fig 3D,E**) suggesting that LE-dependent *NET1* mRNA targeting to the MB is required for NET1 protein function during telophase and abscission. Interestingly, cell accumulation at metaphase was rescued by both the FL 3′UTR and 3′UTR ΔLE transgenes (**Fig S2H**). This indicates that NET1 function during metaphase does not require LE-mediated targeting of *NET1* mRNA, likely due to the disassembly of the nuclear membrane during metaphase and reformation of the nuclear envelope during telophase.

Since inclusion of the *NET1* LE within the 3’UTR is important for NET1 function during telophase, we reasoned that *NET1* mRNA accumulation at the MB may be required for NET1 protein localization to the ICB. It was previously shown that in interphase cells, when cells contain a fully formed nuclear membrane, the majority of NET1 protein is sequestered in the nucleus due to five nuclear localization sequences (NLS) present at the N-terminus of NET1 (Schmidt and Hall 2002; Qin et al. 2005; Alberts and Treisman 1998). It was proposed that a balance between nuclear import and localized cytoplasmic *NET1* mRNA targeting/translation may be needed to regulate NET1 function at specific subcellular compartments (Gasparski et al. 2023; Mason et al. 2025).

To test whether the *NET1* LE is required for NET1 protein targeting to the MB, we stained cells expressing doxycycline-inducible FL 3′UTR or ΔLE 3′UTR transgenes with an anti-HA antibody. We found that in FL 3′UTR expressing cells, HA-NET1 is mostly cytosolic during metaphase and early telophase but accumulates in the ICB during late telophase (**Fig 3F, S3A,B**). We also observed that endogenous NET1 was present in the nucleus as well as in the ICB and MB during late telophase (**Fig S3A**). In contrast, HA-NET1 produced from the ΔLE 3′UTR ΔLE transgene showed significantly less accumulation at the ICB during late telophase (**Fig 3F-G**). Therefore, the efficient accumulation of NET1 protein in the ICB during telophase requires the LE-dependent enrichment of *NET1* mRNA at the same location.

### Localized NET1 mRNA is translated at the intracellular bridge and MB

We and others have shown that specific mRNAs are locally translated at the MB (Farmer et al. 2023; Park et al. 2023). Given that NET1 protein accumulation at the ICB requires LE-dependent *NET1* mRNA targeting to the MB, we then asked if ICB/MB-associated *NET1* mRNA translation may mediate NET1 protein accumulation in the ICB/MB during telophase. To test this hypothesis, we employed a proximity ligation assay (PLA) using antibodies against puromycin and the N-terminal HA tag in our *NET1* rescue transgenes. Puromycin is incorporated into nascent peptide chains during translation and can therefore be used as a marker for newly translated peptides. Since this PLA assay also uses anti-HA antibody, the PLA signal will mark the subcellular locations of nascent HA-NET1 peptides.

In cells expressing the FL 3′UTR *NET1* transgene, we observed puro-PLA signal in the ICB and MB. However, this signal in the ICB was significantly depleted in cells expressing the 3′UTR ΔLE rescue transgene (**Fig 4A,B**), indicating that *NET1* LE is needed to target *NET1* mRNA to the MB and, consequently, to mediate MB-associated NET1 translation. To confirm the specificity of the puro-PLA signal, we included two negative controls: cells without puromycin treatment and cells lacking HA-NET1 expression (**Fig S4A,B**). Both conditions resulted in near-background signal, confirming that the observed puncta were dependent on both the puromycin incorporation in the nascent chain and HA-NET1 transgene expression. From these data, we conclude that *NET1* mRNA is translated at the ICB/MB, but only when the *NET1* mRNA is specifically trafficked to that location.

**Figure 4.**
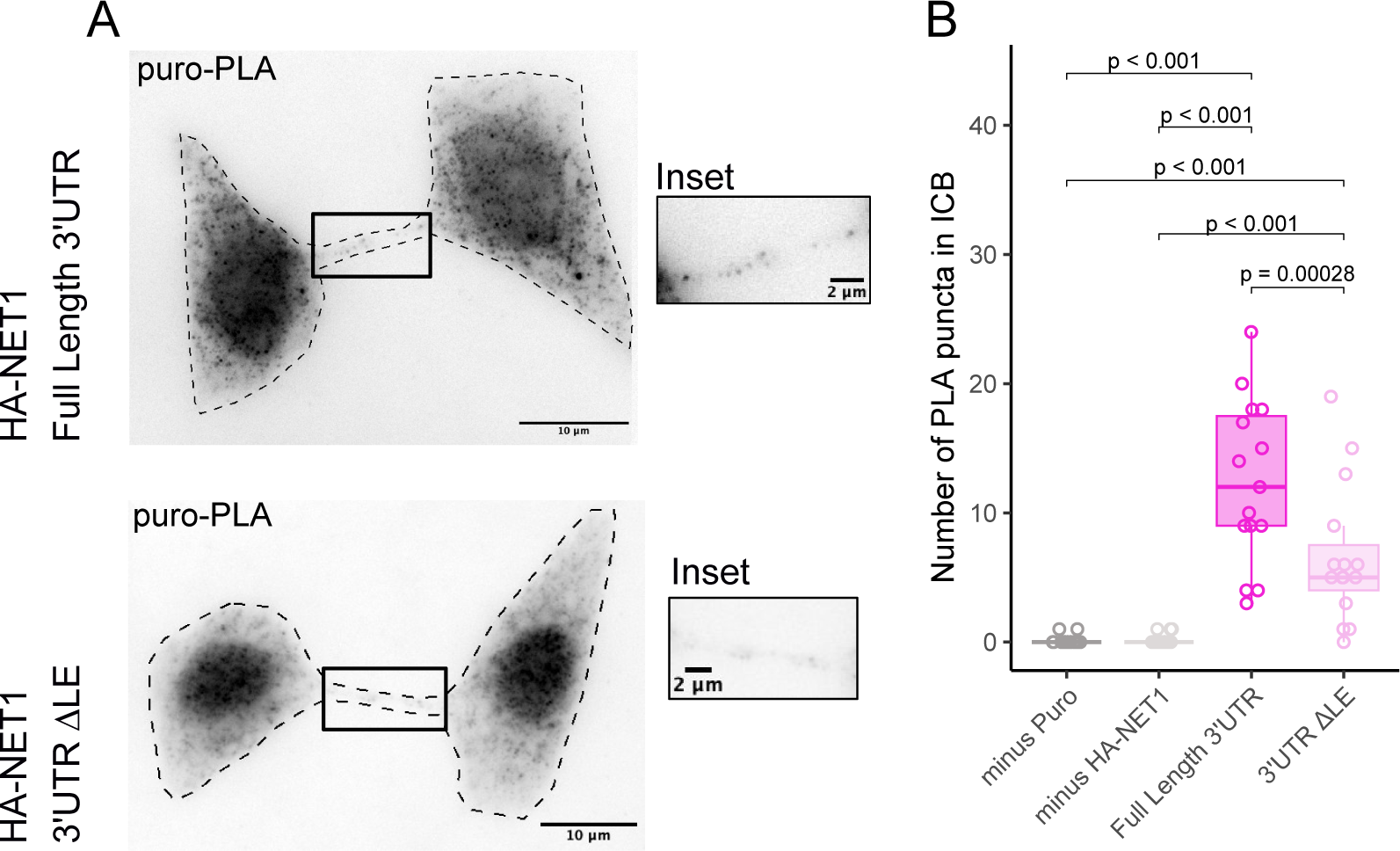
NET1 mRNA is translated at the intracellular bridge and MB. (A) Representative images of a puromycin-proximity ligation assay (puro-PLA, white) images in late telophase cells with antibodies against HA and puromicin. Cell boundaries were defined by GFP-tubulin (not shown). ICB is indicated by the boxed region, and shown in the inset. (B) Quantification of puro-PLA puncta in the ICB across control and experimental conditions P values were calculated using a t-test.

### LE-mediated NET1 mRNA targeting to the ICB/MB is required for Arp2/3-dependent branched actin polymerization at the abscission site

Our data demonstrates that the *NET1* LE drives *NET1* mRNA targeting to the MB and ICB where it is translated and mediates NET1 protein accumulation at the ICB during late telophase. The specific function of localized NET1 during mitotic cell division, though, remained unclear. Inhibiting *NET1* mRNA localization to the ICB led to an increase in ICB length and width, a phenotype repeatedly associated with delayed abscission (**Fig S5A-C**). The same phenotypes were observed in the *NET1* LE PMO-treated cells (**Fig S5D,E**), further implicating ICB-localized NET1 in mediating abscission.

To better understand the role of NET1 during abscission, we performed a time-lapse analysis of live mitotic cells. As it has been suggested that NET1 may function as RhoA GEF (Alberts and Treisman 1998; Schmidt and Hall 2002), we first asked whether the observed defects were due to a loss of RhoA activation during actomyosin contractile furrow formation and ingression. RhoA is activated at the anaphase-telophase transition and is required to form and activate the actomyosin contractile ring (Piekny et al. 2005; Tatsumoto et al. 1999; Kimura et al. 2000; Yüce et al. 2005; Chalamalasetty et al. 2006; Bement et al. 2005). RhoA function during abscission remains unclear, especially as the actomyosin contractile ring needs to be disassembled before abscission occurs (Schiel et al. 2012; Dambournet et al. 2011).

To monitor the spatiotemporal dynamics of RhoA activation during mitosis, we co-expressed a GFP-tagged RhoA biosensor (Mahlandt et al. 2021) and mCherry-α- Tubulin. NET1 KO did not affect RhoA activation during actomyosin ring contractions. Similarly, NET1 KO also did not affect the activation of MyosinIIB, a downstream effector of RhoA, suggesting that RhoA-dependent formation and ingression of the actomyosin contractile ring is not the primary pathway disrupted by NET1 deletion or mis- localization of *NET1* mRNA in mitotic or specifically in late telophase cells (**Fig S6A-D**). These data are consistent with previous studies demonstrating that ECT2, not NET1, is the main RhoA GEF regulating RhoA activity during anaphase and telophase (Kimura et al. 2000; Yüce et al. 2005; Basant and Glotzer 2018; Tatsumoto et al. 1999).

Given that NET1 did not appear to affect RhoA activity at the actomyosin ring, we turned to other GTPases that play important roles in cell division. NET1 has previously been shown to interact with Rac1 (Wang et al. 2024), although it is yet to be determined if NET1 can function as a Rac1 GEF. Further, structural predictions using AlphaFold support the possibility that the NET1 GEF domain may interact with both Rac1 and RhoA (**Fig S6E**).

Arp2/3-induced branched actin mediates the establishment of the abscission site (Advedissian et al. 2024), and cells lacking Arp2/3 accumulation at the abscission site exhibit elongated bridges, consistent with the phenotype observed upon NET1 mis-localization from the ICB. Given that Rac1 activation is also associated with Arp2/3 accumulation (Eden et al. 2002), we asked whether NET1 at the ICB may regulate Rac1-induced Arp2/3-dependent branched actin dynamics during late telophase.

To test this, we used time-lapse microscopy to image the mitotic division of cells co-expressing GFP-Arp3 and mCherry-α-Tubulin. In control cells, we observed GFP-Arp3 accumulation at the abscission site immediately preceding microtubule severing and abscission (**Fig S5G**). GFP-Arp3 accumulation at the abscission site was abolished in NET1 knockout cells and was fully restored by expression of the FL 3’UTR transgene (**Fig 5A,C**). Importantly, expression of the ΔLE 3′UTR transgene failed to fully rescue this defect in Arp3 accumulation at the abscission site (**Fig 5B,C**), further supporting the hypothesis that *NET1* mRNA targeting to and translation at the ICB/MB regulates abscission. All together, these data indicate that the abscission delays observed upon NET1 KO and mislocalization of *NET1* mRNA are a consequence of defects in Arp2/3-dependent branched actin accumulation at the abscission site, thus, delaying the completion of mitotic cell division.

**Figure 5.**
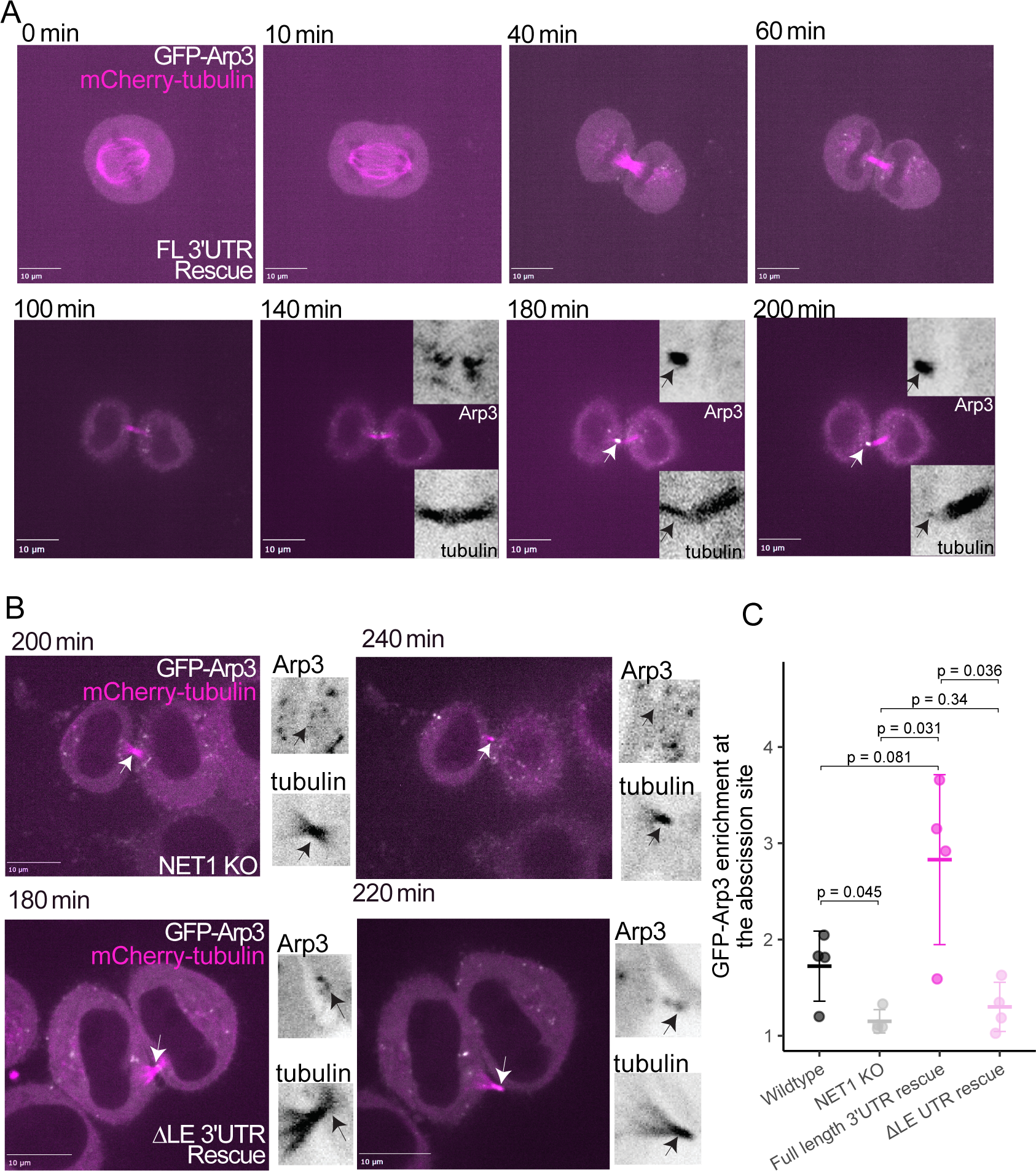
LE-mediated NET1 mRNA targeting to the ICB/MB is required for Arp2/3-dependent branched actin polymerization at the abscission site. (A) Time-lapse live cell imaging of NET1 KO cells with HA-NET1 FL 3’UTR rescue. Cells expressing GFP-Arp3 (white) and mCherry-tubulin (magenta) were imaged from metaphase through abscission. Abscission is indicated by the loss of tubulin on one side of the ICB. Insets show ICB Arp3 accumulation in late telophase preceding abscission. (B) Telophase cells expressing GFP-Arp3 (white) and mCherry-tubulin (magenta). Cells immediately before and after abscission are shown with insets for Arp-3 and tubulin at the ICB. (C) Quantification of Arp-3 expression at the abscission site relative to total ICB Arp-3 expression. P values were calculated using a t-test.

### Arp2/3 accumulation at the midbody requires NET1 binding to Rho-family GTPases

We have established that LE-dependent localization of *NET1* mRNA is required for Arp2/3-mediated branched actin accumulation at the abscission site in late telophase. Since NET1 is a GEF for Rho family GTPases, we hypothesized that NET1 functions by mediating Rac1 activation at the abscission site. We therefore asked if NET1 interaction with GTPases via its GEF domain is required for its function in lat telophase. To test this, we took advantage of a previously reported point mutation in NET1, L321, that blocks NET1 binding to RhoA and likely to Rac1 (**Fig S6E**), presumably rendering NET1 catalytically inactive (Alberts and Treisman 1998; Menon et al. 2013; Rossman et al. 2005; Schmidt and Hall 2002).

We rescued our NET1-KO cell line with a transgene encoding HA-NET1-L321E-FL3’UTR (**Fig 6A**). This transgene contained the full length *NET1* 3’UTR and was therefore efficiently targeted to the ICB (**Fig S7B,C**) but was unable to interact with Rho family GTPases via its GEF domain due to the L321E point mutation (Alberts and Treisman 1998). Importantly, the NET1 binding mutant was expressed at similar levels to the other HA- NET1 rescues (**Fig S7D**). We then analyzed Arp2/3 accumulation at the abscission site in these cells using time-lapse microscopy. The HA- NET1-L321-FL3’UTR construct did not rescue GFP-Arp3 accumulation at the abscission site (**Fig 6C**), indicating that GEF domain binding is needed for NET1 function during abscission. Cells expressing the NET1 binding mutant also exhibited significantly elongated intercellular bridges, a hallmark of delayed abscission, further suggesting that this mutant was unable to rescue abscission defects **(Fig S7A)**. The localization of NET1 mRNA and protein to the midbody is therefore not, on its own, sufficient to promote timely abscission. The ability of localized NET1 protein to interact with GTPases is also essential.

**Figure 6.**
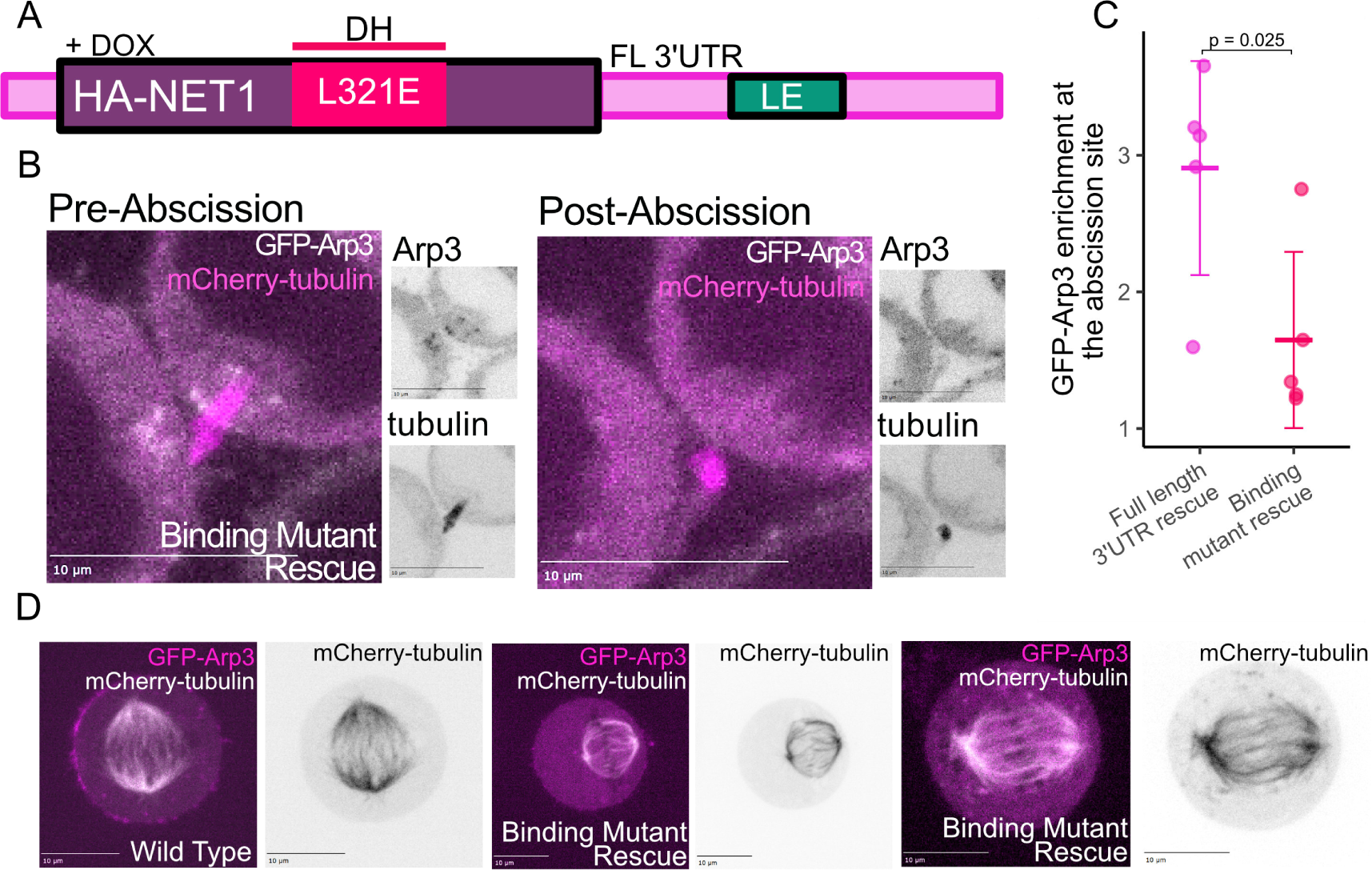
Arp2/3 accumulation at the midbody requires NET1 binding to and Rho-family GTPases. (A) Schematic of HA-NET1 rescue construct indicating the location of the amino acid substitution resulting in a GTPase binding mutant. (B) Telophase cells expressing GFP-Arp3 (white) and mCherry-tubulin (magenta). Cells immediately before and after abscission are shown, with insets for Arp3 and tubulin at the ICB. (C) Quantification of Arp3 expression at the abscission site relative to total ICB Arp-3 expression. P values were calculated using a t-test. (D) Representative images of wildtype HeLa cells and cells and NET1 knockout cells expressing HA- NET1-L321E in metaphase. Spindle microtubules are labeled with anti-acetylated tubulin antibodies (white). Images show the defects in mitotic spindle assembly (middle image) and positioning (right image).

Interestingly, expression of the NET1-L321 mutant also affected mitotic spindle assembly and positioning (**Fig 6D**), suggesting that GEF domain binding is also required for NET1 function during metaphase. While our system is based on complete loss of endogenous NET1, others have shown that cells treated with *NET1* siRNA and rescued with an siRNA-resistant NET1 point mutant exhibit normal mitotic spindles (Menon et al. 2013). A small amount of wildtype NET1 expression may therefore be sufficient to rescue this defect.

Nevertheless, the reason behind these differences remains unclear, and further studies will be needed to better understand how NET1 regulates mitotic spindle assembly.

Together, our data demonstrates that faithful completion of abscission in late telophase cells requires both NET1 localization to the ICB and interaction of the NET1 with its GTPase effectors, likely Rac1, to drive Arp2/3- mediated branched actin accumulation at the abscission site for a faithful and timely competition of mitosis (**Fig 7**). These results establish the mechanism behind, and importance of, the localization of a specific RNA to the MB and serve as a framework for the investigation into other RNAs that may similarly regulate the process.

**Figure 7.**
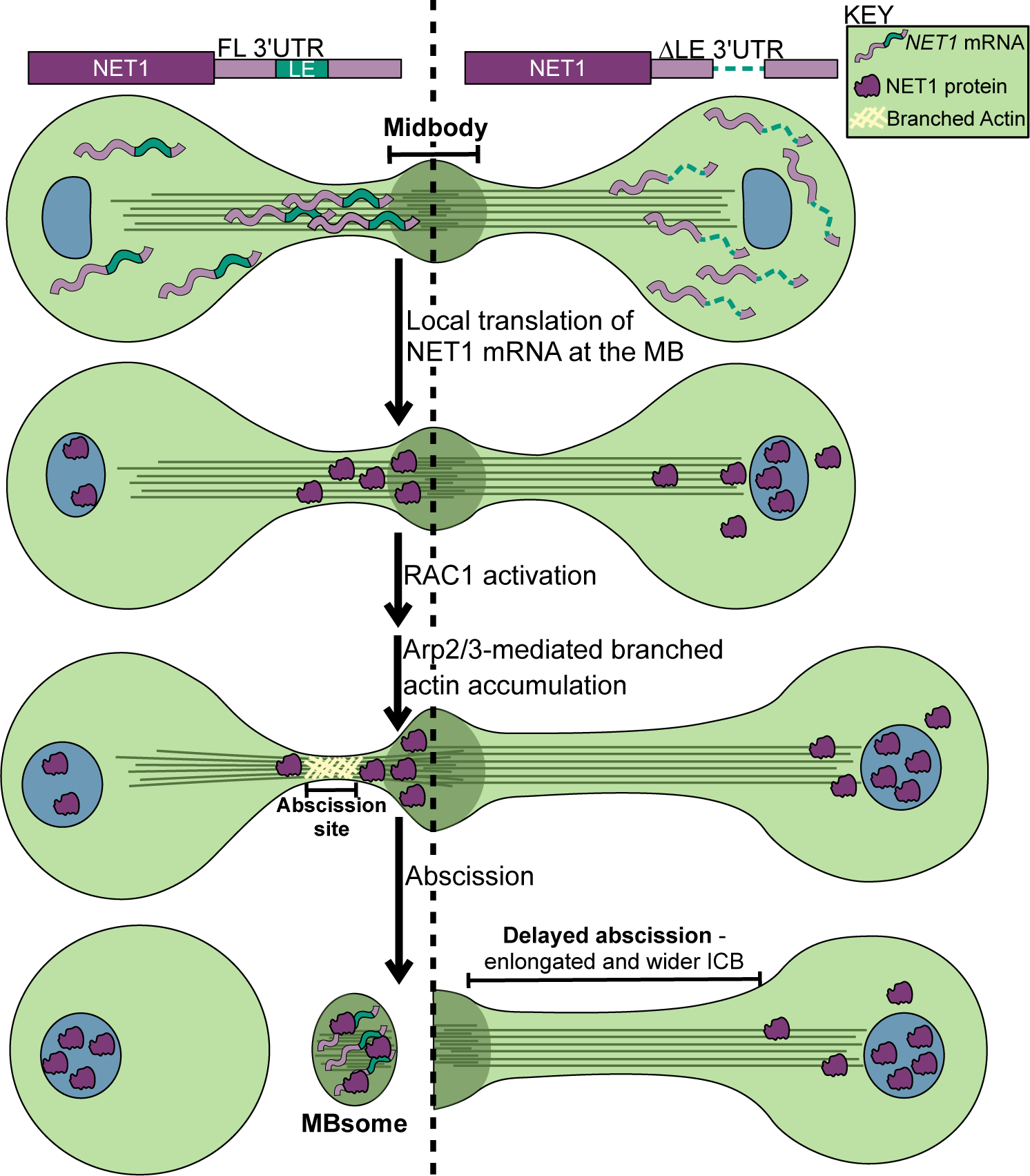
Model for *NET1* mRNA trafficking and its role in Arp2/3-dependent abscission. NET1 mRNA localization to the midbody is dependent on a localization element within its 3’UTR. MB localized NET1 mRNA is translated in the ICB, resulting in local NET1 protein enrichment at the MB. NET1 GEF activity in the ICB, likely via Rac1 activation, results in Arp2/3 accumulation at the abscission site to enable timely abscission. When NET1 mRNA is lost from the MB, either through NET1 knockout or mis-localization, Arp2/3 fails to accumulate at the abscission site, and abscission is delayed.

## DISCUSSION

Mitotic abscission is a complex process that involves coordination of membrane fusion, microtubule remodeling and severing, and localized dynamic changes in the actin cytoskeleton. All of these processes need to be precisely timed and executed during abscission. However, despite the importance of abscission for the completion of mitotic cell division, the molecular machinery governing this process remains to be fully understood.

Major open questions include how the location and timing of abscission is determined and how the proteins mediating abscission are targeted to the midbody. Recent studies have demonstrated that the midbody contains a set of specific mRNAs (Farmer et al. 2023; Park et al. 2023). Interestingly, these mRNAs encode many well-established abscission regulators, such as MKLP1, all ESCRT-III complex subunits, and Citron Kinase (Farmer et al. 2023; Park et al. 2023). Furthermore, the MB proteome contains the machinery necessary for protein translation (Farmer et al. 2023; Addi et al. 2020; Park et al. 2023). Thus, it was proposed that mRNA localization and localized translation at the MB and ICB may contribute to targeting and assembly of necessary abscission machinery at the abscission site (Farmer et al. 2023). What remained unknown, however, are the mechanisms mediating mRNA localization to the MB and whether targeting of these specific mRNAs to the MB plays a role in regulating abscission.

In this study, we identify mechanisms that regulate *NET1* mRNA localization to the MB and its local translation. Using a massively parallel reporter assay, we identified a LE within the 3’UTR of *NET1* mRNA that was necessary and sufficient for transport to the MB. This region overlapped strongly with a region of the 3’UTR previously shown to result in trafficking of the mRNA to the basal pole of epithelial cells where it contributed to the organization of the interface between the epithelial cells and the basement membrane (Mason et al. 2025). Given that both the MB and the epithelial basal pole lie at the plus-end of microtubules, this lends further support to a previously suggested model that mRNAs share a core machinery responsible for microtubule-based targeting that operates across many different cell types (Goering et al. 2023).

Determining the LE driving *NET1* mRNA localization to the MB is an important step in understanding the mechanism responsible for MB RNA localization, but it does not on its own demonstrate the importance of *NET1* mRNA localization for cell division and abscission. Indeed, there are relatively few reported examples demonstrating that the mislocalization of a specific RNA in mammalian cells leads to defects in cellular function. However, the identification of the LE within the mRNA put us in position to ask this question as it gave us a specific sequence to target in order to perturb *NET1* mRNA localization. Using both LE-targeted PMO and knockout/rescue approaches, we found that perturbation of *NET1* mRNA localization decreases cell proliferation rates and does so at least in part through inhibiting abscission.

NET1 has five nuclear localization sequences (NLS) present at the N-terminus of the protein (Schmidt and Hall 2002; Qin et al. 2005). As a result, NET1 is rapidly transported into the nucleus where it is sequestered from functioning in the cytoplasm (Schmidt and Hall 2002). We propose that during telophase, after reformation of the nuclear membrane, NET1 protein accumulates at the abscission site through local MB-associated translation of *NET1* mRNA. This local translation directly at the ICB may be important for the ability of MB-localized NET1 to escape the transport to the nucleus that would otherwise happen due its NLS sequences (Gasparski et al. 2023; Mason et al. 2025).

This hypothesis also explains why metaphase defects can be rescued by both the HA-NET1 ΔLE 3’UTR and the HA-NET1 FL 3’UTR rescue. During metaphase, cells break-down the nuclear envelope. Local translation is therefore not needed during metaphase to keep NET1 from being sequestered in the nucleus. These data identify mRNA localization and local translation at the MB/ICB as a key regulatory layer mediating spatiotemporal dynamics of abscission machinery during late telophase.

The loss of MB-localized *NET1* mRNA, and therefore MB-localized NET1 protein, results in telophase defects, implicating NET1 as an important regulator of the abscission. What remained unclear, however, is how NET1 actually functions during abscission. NET1 is a Rac/Rho guanine nucleotide exchange factor (GEF) that often activates RhoA, a small monomeric GTPase, during cell migration (Qin et al. 2005). Surprisingly, we found that NET1 likely plays a role outside of RhoA activation to facilitate abscission. We instead found that, during late telophase, NET1 may act as a regulator of Rac1. AlphaFold modeling suggests NET1’s GEF domain interacts similarly with both Rac1 and RhoA, and previous studies have shown NET1 binding and interaction with Rac1, suggesting that NET1 may act as a Rac1 GEF (Carr et al. 2013; Wang et al. 2024).

Localized Arp2/3-induced branched actin polymerization determines the site and timing of abscission, presumably by regulating ESCRT-III polymer formation and spastin-dependent microtubule severing (Advedissian et al. 2024). However, it is unknown what induces and regulates *de novo* Arp2/3 activation at the abscission site. Rac1 is known to activate Arp2/3-dependent branched actin polymerization and our data suggests that NET1 may activate Rac1. We found that loss of NET1 in the ICB either through knockout or mis- localization of *NET1* mRNA resulted in a loss of Arp2/3 mediated branched actin accumulation at the abscission site. Taken together, these findings suggest that localized translation of *NET1* RNA at the abscission site may be a key step in determining the location and timing of the abscission site.

To further test whether NET1 GEF activity is required for branched actin accumulation, we used a NET1 point mutant, L321E, in our knockout/rescue assays. The L321E mutation is within the DH domain and has been shown to disrupt NET1 binding to Rho GTPases, thereby likely rendering NET1 catalytically inactive (Alberts and Treisman 1998; Menon et al. 2013). Consistent with the requirement of NET1 GEF activity during abscission, the expression of this mutant NET1 did not rescue Arp2/3 accumulation at the abscission site during late telophase. Thus, the targeting and local translation of *NET1* mRNA is required for the regulation of Arp2/3-dependent branched actin accumulation at the abscission site, presumably via activation of Rac1.

The finding that NET1 may regulate Rac1 activation came as a surprise since originally NET1 was identified as a RhoA GEF. There are two possibilities explaining this conundrum. The first possibility is that NET1 may be a GEF for both RhoA and Rac1. Indeed, GEFs are known to be promiscuous and often the specificity of their GEF activity is determined by other factors, such as the localized presence of a specific small monomeric GTPase. Furthermore, post-translational modifications can also affect GEF activity. For example, it has been shown that phosphorylation of NET1 by Cdk1 during metaphase inhibits its GEF activity toward RhoA (Ulu et al. 2021). The second possibility is that NET1 may regulate Rac1 not as a GEF but rather as a scaffolding protein. Indeed, it was recently shown that NET1 binding to Rac1 can inhibit its degradation (Carr et al. 2013). Whether during mitosis NET1 functions as Rac1 GEF or if it regulates Rac1 activity indirectly remains to be determined.

In this study, we have identified a key regulatory mechanism that mediates faithful progression through mitosis and abscission via *NET1* mRNA localization and translation at the MB and ICB. This adds to the small but growing number of known roles for the localization of specific mRNAs in regulating key mammalian cellular processes. We also identified NET1 as an important regulator of abscission and have shown that in late telophase NET1 contributes to the establishment of the abscission site by facilitating localized Arp2/3 and branched actin accumulation. While this study defines a new mode of regulating abscission, many questions remain. These include the identities of the kinesins and RNA-binding proteins that mediate *NET1* mRNA transport to the MB, the mechanisms that regulate the timing of the translation of these mRNAs, and whether mRNA transport and local translation are regulated by the abscission checkpoint. Further studies will be needed to answer all these questions.

## STUDY LIMITATIONS

This study suggests that mRNA targeting and translation at the MB/ICB is needed for abscission. However, most of the data suggesting local, MB-associated, translation is correlational. At this point, we do not have a way of inducing a localized inhibition of translation specifically at the MB. Thus, we cannot directly test whether MB-associated translation is required for completion of mitotic cell division. Additionally, all of the experiments in this study have been done in HeLa cells. It therefore remains to be determined whether the observed defects in abscission upon mistargeting *NET1* mRNA is a peculiarity of cancer-derived cells or is a wider phenomenon in many cell types and organisms.

## ACKNOWLEDGEMENTS

We are thankful to Drs. Stavroula Mili, Devon E. Mason, and Konstadinos Moissoglu for insightful discussions about NET1 mRNA targeting and for providing us with PMOs. This work was funded by R35-GM133385 (to JMT), F31-GM155957 (to KV), R01-GM143774 (to RP), and P-MIP-25-115 (to RP).

## DATA AVAILABILITY

Raw data associated with the MPRA can be accessed at the Gene Expression Omnibus using accession number GSE342282.

## METHODS

### Cell Culture

HeLa cells were grown in DMEM supplemented with 10% Fetal Reserve Bovine Serum and 1% penicillin– streptomycin solution. The cells were grown in a humidified incubator at 37°C and 5% CO_2_. Cells were regularly screened for mycoplasma and found to be negative.

### Midbody Purification from Media and RNA isolation

Midbodies were isolated following this protocol (Peterman and Prekeris 2017). Hela cells were grown in 1 µg/ mL doxycycline to induce NET1 rescue or reporter expression for 48 hours. Cells were grown to 90% confluent in 24x 15cm plates over the 48 hours. All subsequent steps are performed at 4°C. Conditioned media was collected from cells and centrifuged at 3,000xg for 5 minutes twice to pellet cells and cellular debris. The supernatant was pooled and centrifuged at 10,000xg for 30minutes. The pellet was washed in 1mL PBS and again centrifuged at 10,000xg for 30 minutes. This pellet was resuspended in nuclease-free water, 6 mM MgCl2, 250 mM sucrose, and 250 mM NaCl to a final volume of 200 µL. In a 1.3mL ultracentrifuge tube, a step-density gradient was created consisting of 400 µL 1M sucrose and 400µL of 2M sucrose on top of a 300µL 40% glycerol cushion. Sucrose gradients are made in a buffer of 20 mM HEPES, pH 7.4, 6 mM MgCl2, 1 mM Dithiothreitol, and 100 mM NaCl. The resuspended pellet was layered on top of the sucrose step gradient and centrifuged in a TLS-55 Swinging-Bucket Rotor at 3,000xg for 30 minutes. The band between the 40% glycerol and 2M sucrose interphase layers is collected and pelleted at 10,000xg for 45 minutes in a fresh 1.5mL centrifuge tube. The supernatant is discarded and the MB pellet is retained for RNA isolation using a Zymo Quick-RNA Microprep Kit. A whole cell sample for comparison is isolated by trypsinising a subset of the cells that produced the conditioned media for RNA isolation using a Zymo Quick-RNA Miniprep Kit.

### Oligo nucleotide design for MPRA

The code for designing the MPRA oligos is available at LINK. The script created 260nt oligonucleotides with a step size of 4nt between neighboring oligonucleotides. Oligonucleotides were designed against the *NET1* 3’UTR. This transcript contained well-defined 3’UTR ends and 260nt were added upstream and downstream of the 3’UTR start and end to give full coverage of the ends of the 3’UTR with multiple oligonucleotides. 20nt PCR handles were added to the ends of every oligonucleotide. The pool of oligonucleotides was synthesized by Twist Biosciences.

### Oligonucleotide library cloning

The oligonucleotide pool obtained from Twist Bioscience was resuspended in 10mM Tris-EDTA buffer, pH 8.0 to a concentration of 10 ng/µL. The pool was amplified by performing 8x 50µL PCR reactions. Each reaction contained an input DNA template of 10ng from the original pool. The PCR reaction used KOD hot Start Master Mix with 15 amplification cycles. Following amplification, the PCr reaction was treated with Exonuclease I at 37°C for 2 hours to digest the single-stranded template and primers. The DNA was then purified using the NucleoMag NGS Clean-up and Size Select protocol. In the first purification round, 0.6× NucleoMag beads were used to get rid of longer DNA products. The supernatant from this purification was then removed, and additional NUcleoMag beads were added to bring the final overall concentration to 1×. DNA bound to these beads was then washed and eluted in TE buffer.

The pRD-RIPE plasmid was linearized by digestion with BstXI at 37°C for 4hr to clone the oligonucleotide library into the 3’UTR of the GFP reporter. Digested plasmid DNA was gel extracted using Zymoclean Gel DNA Recovery Kit. The digested plasmid and amplified DNA library were ligated using Gibson Assembly reaction using the insert:vector ratio molar ratio of 7:1 at 50°C for 1hr. The cloned reporter plasmid (∼200ng) was ethanol precipitated to get rid of excess salts and then was transformed into NEB® 10-beta Competent *E. coli* using a Biorad GenePulser electroporator. The transformed cells were grown in the recovery medium supplied with the cells at 37°C for an hour and then added to 500mL Luria broth (LB) in a 1L flask and incubated overnight at 37°C shaking at 250rpm. The next day the bacterial culture was centrifuged at 4000rpm for 20min. The reporter plasmid libraries were purified using ZymoPURE Plasmid Maxiprep kit. Restriction digestion was performed to confirm that the plasmid library contains only a single insert of the right size.

### Targeted RNA sequencing of MPRA library

1000ng total RNA from each MB and WC sample was input into a 20µL reaction using SuperScript IV Reverse Transcriptase to generate cDNA. The reaction was performed according to the manufacturer’s protocol with primers specific to the GFP CDS containing an 8-nt unique molecular identifier(UMI) and a partial Illumina read 1 primer sequence. The incubation time at extension step (55°C) was increased to an hour. Post reverse transcription, 1µL each of RNAse H and RNAseA/T1 mix was added directly into the RT-reaction and incubated at 37°C for 30min to digest the remaining RNA and RNA:DNA hybrids. The cDNA was purified using Zymo DNA Clean & Concentrator kit using 7:1 excess of DNA-binding buffer recommended for binding ssDNA. For library preparation, purified reporter cDNA reaction was split into 5 PCR reactions (4µL cDNA/PCR) and amplified using a reporter specific forward primer with Illumina sequencing adaptors using KOD hot Start Master Mix using 24x cycles. The five PCR reactions per sample were pooled together and purified using NucleoMag NGS Clean-up and Size Select protocol. In the first purification round, 0.6× NucleoMag beads were used to get rid of longer DNA products. The supernatant from this purification was then removed, and additional NUcleoMag beads were added to bring the final overall concentration to 1×. DNA bound to these beads was then washed and eluted in TE buffer. The library was quantified using Qubit dsDNA HS Assay Kits and the size of the library was verified using Tapestation.

### Analysis of MPRA results

Adaptors were removed from reads using cutadapt (Martin 2011). Specifically the sequences GGCGGAAAGATCGCCGTGTAAGTTTGCTTCGATATCCGCATGCTA and CTGATCAGCGGGTTTCACTAGTGCGACCGCAAGAG were trimmed from the 5’ends of the forward and reverse reads respectively. The trimmed reads were then aligned to the reference oligonucleotide sequences using bowtie2 and the following parameters: -q –end-to-end –fr –no-discordant –no-unal -p 4 -x Bowtie2Index/ index -1 forreads.fastq -2 revreads.fastq -S sample.sam. Typically, 99% of reads had the expected adapters, and 95% of those aligned to the reference oligonucleotides.

The number of unique UMIs (the first 8 nt of the reverse read) for each reference oligonucleotide was then calculated using https://github.com/TaliaferroLab/OligoPools/blob/master/analyzeresults/UMIsperOligo.py. These UMI counts were then given to DESeq2 (Love et al. 2014) to quantify oligonucleotide abundances in each sample and identify oligonucleotides enriched in MB or WC samples.

### Assaying MB RNA localization via RT-qPCR

Purified RNA from MB and WC samples was isolated as previously described. 100ng of RNA per sample was reverse transcribed in a 10µL reaction using LunaScript RT Supermix per manufacturer’s instructions. The cDNA was diluted to 20µL total in nuclease-free water. For each qPCR reaction, 2 µL of diluted cDNA is used as the template. For the reporter assay, qPCR was used to determine the abundance of Firefly and Renilla reporter transcripts in the MB and WC samples. For rescue assay, qPCR was used to determine the abundance of NET1 and TSG101 transcripts in the MB and WC samples. Using PrimeTime® Gene Expression Master Mix the qPCR reaction was performed using probe sets with different labels so that each pair of transcripts would be quantified in the same qPCR reaction. Firefly and TSG101 are HEX labeled. Renilla and NET1 are FAM labeled.

Reactions were performed using CFZ-Opus 384 thermocycler under these conditions: UNG activation at 50 °C for 2 min, followed by polymerase activation at 95 °C for 30s and 40 cycles of 95 °C for 5s, and 60 °C for 30s. A melting curve was performed by incubating samples at 65°C for 15s followed by a temperature gradient increase at 0.5°C/s to 95°C. Each sample was quantified in three technical replicates. Further, no RT and no template controls were performed. Fold enrichment was quantified using ΔΔCt method. MIQE guidelines were followed for all qPCR experiments.

### Single-molecule RNA FISH probe design

The smiFISH protocol was adopted from (Tsanov et al. 2016). Briefly, the probes were designed by inputting the gene sequences (NET1, and TSG101) into the Oligostan software. smiFISH probes then had a Y flap (5′- TTACACTCGGACCTCGTCGACATGCATT-3′) appended such that we could hybridize the fluorescent molecules to the probe. The fluorescent probe was designed by taking the reverse complement of the Y flap (5′-AATGCATGTCGACGAGGTCCGAGTGTAA-3′) and adding Cy3 to both ends. smFISH probes for Firefly luciferase were obtained from Biosearch Technologies. The probes were labeled with Quasar 570 dye. The sequences of these probes are proprietary.

### Visualization of RNA localization using single-molecule RNA FISH

HeLa cells were plated on PDL-coated coverslips and induced with 1 µg/mL doxycycline for 48hrs, grown to a maximum of 50% confluent. The media was aspirated and cells were washed once with 1× PBS. For Halo-tag visualization, HaloTag Oregon Green Ligand was added 4–6 h prior to fixing cells in the incubator. Cells were fixed in 3.7% formaldehyde for 10 min at room temperature and then washed twice with 1× PBS. Cells were permeabilized with 70% ethanol at 4°C for 2hrs. The cells were washed with freshly prepared wash buffer (2X SSC and 10% formamide in water) at room temperature for 15 min. In the meantime, the smiFISH probes for each gene were hybridized to the fluorescent Y Flap using the protocol from Tsanov et al, 2016 (Tsanov et al, 2016). NET1 and TSG101 probe sets each included 48 probes.

After hybridization, the probe/flap hybridization product was spun in a benchtop centrifuge for 60 s. Per coverslip, 2 µl (0.833 µM) of probe was added to 100 µl of smFISH hybridization buffer activated with 10% formamide for each condition and gene. A hybridization chamber was prepared using an empty tip container, wrapped in tinfoil with parafilm and wet paper towels inside the box to retain moisture. 100 µl of the probe-containing hybridization solution was added to the parafilm. The coverslip was then placed on top of this droplet of hybridization buffer with the cell side down. The hybridization chamber with the coverslips was incubated at 37 °C overnight (15–18 h). The coverslips were transferred to a fresh 12-well plate with the cell side up and incubated twice with freshly prepared wash buffer for 45 min at 37 °C. Then, the slides were incubated with wash buffer including DAPI (100 ng/mL) at 37 °C for 30 min. Slides were washed twice with PBS for 5 min at room temperature. Coverslips were then mounted onto slides with Fluoromount G and sealed with nail polish.

Slides were imaged with a 63x oil immersion lens with consistent laser intensity and exposure times across samples. DAPI was imaged with an exposure of 10 ms. The FITC channel was imaged with an exposure of 400 ms. FISH probes were visualized in the TRITC channel with an exposure of 1500 ms. Z stacks were collected of ∼24 images, 0.4 µm apart.

### Analysis of single-molecule RNA FISH data

FISH-quant was used to quantify midbody or intercellular bridge enrichment of smFISH spots as previously described(Tsanov et al. 2016; Farmer et al. 2023). Briefly, outlines were drawn in the FITC channel visualizing midbody via Halo-MKLP1 fluorescence or the intercellular bridge via GFP-tubulin fluorescence. Three outlines were drawn per dividing cell: two cells on each side of the bridge, and the intercellular bridge. Prior to quantification, identified smFISH spots were thresholded for intensity, sphericity, amplitude, and position. RNA-FISH enrichment was quantified by the total number of spots in the intercellular bridge of cells over a total number of spots in the whole cell normalized to the control.

### PMO delivery

Cells were grown to 75% confluence in 12 well plates and the media was replaced with 750µL of fresh media. Cells were then transfected with the indicated PMO (NET1 LE, NET1 3’UTR control or scramble) at a concentration of 15 µM (15µL of PMO). Media was swirled gently and then 4.5µL of Endoporter was added and media was immediately swirled gently. Cells were treated overnight and the media was changed the next morning (15-18hrs). For imaging experiments, cells were split into 5x 12 wells per PMO transfection containing PDL-coated coverslips. 48-72 hours after transfection cells were fixed and stained as described in RNA-FISH or Immunofluorescence method. For quantification of NET1 RNA levels, RNA was harvested from PMO treated whole cells and used as input for qPCR method as described. NET1 levels were normalized to control (TSG101) levels in each sample and compared to a non-PMO treated control.

### Cell Proliferation Assay

Cells were seeded at a density 0.01 x 10_6_ of cells per well. For PMO treated cells, cells were seeded the morning after PMO transfection. For dox inducible rescues, cells were seeded in media containing 1 µg/mL doxycycline. Plates are maintained in the Incucyte® over the duration of the proliferation assay in an incubator at 37°C and 5% CO_2_. 4 phase contrast images at 10x magnification were taken per well every 3 hours for 72-96hours. Cell confluency was measured using Sartorius software for each timepoint and well over the timecourse.

### Immunofluorescence and Quantification

HeLa cells were treated as indicated in the text and then fixed in 1% formaldehyde in PBS for 10 min at room temperature. Cells were rinsed three times in PBS. The cells were then incubated with a primary antibody in PBS containing 0.5% BSA and 0.2% saponin for 1 h at room temperature, washed three times in PBS, and then incubated with the appropriate fluorochrome-conjugated secondary antibodies diluted in PBS containing 0.5% BSA and 0.2% saponin for 30 min. Cells were washed three times in PBS and mounted with Fluoromount.

Fixed cells were imaged with an inverted Axiovert 200M microscope (Zeiss) with a ×63 oil immersion lens and QE charge-coupled device camera (Sensicam). Z-stack images were taken at a step size of 500–1,000 nm.

Image processing and quantification were performed using 3i Slidebook 6 software (Intelligent Imaging Innovations). Briefly, masks (ROI) were made to calculate the total intensity of the cell body or the intercellular bridge as well as the total area for each mask. The numbers were then used to calculate the intensity/µm2. Unless otherwise stated, all images are represented as a maximal projection of the z-stack.

### Creation of NET1 knockout cell lines with a single loxP cassette

The guide RNA (gRNA) sequences AAGGACGUGACACGAGCCAG and ACGUGUUCCACAAGCAGCAG for *NET1* were obtained from synthego and dissolved in 1x TE buffer for a final concentration of 100µM. Purified Cas9 protein was added to the teo gRNAs at a 3:1 molar ratio of gRNA:Cas9 at room temperature for 15 minutes. Then, these RNP complexes were co-electroporated with a CMV-GFP plasmid into Hela cells using the Neon transfection system with the following settings: 2 pulses at 1005V for 35ms. 48 hours after transfection, cells were sorted into single GFP positive cells with a flow cytometer and allowed to grow for two weeks. Clones were screened for NET1 expression by polyclonal NET1 antibody and RNA sequencing. Loss of the NET1 locus or lesions at the NET1 locus was identified by PCR from genomic DNA from single cell clones.

### Expression of NET1 rescue constructs in Hela Cells

Hela NET1 KO cells were plated in a 6 well plate at approximately 6 × 10^5^ per well in DMEM media supplemented with 10% Fetal Reserve for 48 hours before transfection. Cells were co-transfected with 1% wt/ wt of a Cre encoding plasmid (pCAGGS-Cre) and one of the plasmid derivatives containing dox-inducible HA- NET1 rescue coding sequence with either the full length *NET1* 3’UTR or the *NET1* 3’UTR with the localization element (defined by the MPRA) deleted. For the transfection of one well of a 6 well plate, 0.5 µg of NET1- rescue plasmid was mixed with 20 ng of Cre-encoding plasmid, 8 µL of Lipofectamine LTX, 4 µL PLUS reagent and 200 µL Optimem based on the manufacturer’s protocol. Cells were incubated overnight with the transfection reagents before the media was replaced and cells were incubated for an additional 24 hours. 5.0 µg/mL puromycin was added until cells in control wells were fully depleted. Remaining cells with stably integrated rescue constructs were expanded in full growth media plus puromycin. Lysates were then collected to assess rescue construct expression by western blot and immunofluorescence.

### Proximity ligation assay

NET1 KO cells were transfected with either HA-NET1 FL 3’UTR or HA-NET1 ΔLE 3’UTR constructs. Non-transfected NET1 KO cells were used as a negative control. The other negative control used was a non-puromycin treated cells transfected with HA-NET1 FL 3’UTR. Cells were then treated with 2µg/mL puromycin for 10 min, followed by fixation with 4% paraformaldehyde and permeabilization with 0.1% Triton X-100. Cells were incubated with anti-HA and anti-puromycin antibodies, followed by Duallink proximal ligation assay (PLA), as described in the manufacturers protocol (Sigma-Aldrich). For imaging, cells in late telophase were randomly picked using the following criteria: (1) cells had fully reformed round nuclei; (2) cell bodies were flattened; (3) daughter cells were connected by an extended intercellular bridge of at least 2 µm in length. For quantification of ICb puncta, we followed this protocol (De Pace et al. 2025). Briefly, the images were max projection, and channels separated. Using the brightfield channel the ICB was selected. From this selection in the Puro-PLA channel, the signal was thresholded to identify spots of correct size (filtering for spots larger than 20 µm^2^, anything smaller is likely background signal), finally using the same thresholding for all images the number of spots were counted.

### Immunoblotting

For characterization of plus/minus HA-NET1 expression, cells were incubated with or without doxycycline (2.0µg/mL) for 48 hours. Cellular lysates were collected in 50 µL of ice cold RIPA buffer, incubated on ice with periodic vortexing for five minutes before centrifugation at 21000 rcf for 5 minutes. The supernatant was removed and 15µL of the lysate was added to sample buffer and DTT at a final concentration of 3 mM and boiled at 95°C for five minutes. Remaining lysate was stored at −80°C. Denatured lysates were separated by PAGE on 4%−12% Bis-Tris gradient along with Spectra Protein Ladder in MOPS SDS Running Buffer at 150V for 90 minutes. The gel was transferred to a PVDF membrane using an iBlot2 dry transfer. Total protein was assessed with Ponceau staining and destained before blots were blocked while rocking in 5% milk powder in PBST for 45 minutes at room temperature. Blots were subsequently washed 3 times for 5 minutes each in PBST at room temperature before incubation with respective antibody in PBST overnight at 4°C with agitation. Blots were washed three times for 5 minutes each with agitation and incubated with their respective HRP-conjugated secondary antibody for 30 minutes at room temperature. Blots were washed 3 times for 5 minutes in PBST and visualized with the WesternBright Sirius HRP kit on a Sapphire imager set to collect chemiluminescence signal.

### Microscopy analysis

#### Immunofluorescence microscopy

HeLa cells were treated as indicated in the text and then fixed in 1% formaldehyde in PBS for 10 min at room temperature. Cells were rinsed three times in PBS. The cells were then incubated with a primary antibody in PBS containing 0.5% BSA and 0.2% saponin for 1 h at room temperature, washed three times in PBS, and then incubated with the appropriate fluorochrome-conjugated secondary antibodies diluted in PBS containing 0.5% BSA and 0.2% saponin for 30 min. Cells were washed three times in PBS and mounted with Fluoromount. All fixed cell imaging was performed on a widefield inverted Zeiss Axiovert 200M microscope using a 63X oil-objective, QE charge-coupled device camera (Sensicam), and Slidebook v. 6.0 software (Intelligent Imaging Innovations). Images were taken as z-stacks with 0.5 µm step intervals. Where indicated images were deconvolved (Nearest Neighbors) using the Intelligent Imaging Innovations software. For ICB morphology analysis images were max projected to include all stacks (200nm step size) in which the central spindle microtubules (as labeled with anti-acetylated tubulin antibodies) was in focus. The lengths and width (at the narrowest part) of central spindle microtubules from the resulting projection was then measured. Only cells in late telophase were analyzed. The main criteria to classify mitotic cells in late telophase was used as previously described (Sachs et al. 2026). Ten to fifteen cells were analyzed from each independent biological replicate and experiment was repeated three times.

#### Time-lapse microscopy and analysis

For time-lapse analysis Hela cells were co-transfected with mCherry-tubulin and GFP tagged Myosin IIB, Arp3, or RhoA bisoensor and imaging was performed using Zeiss Axio Observer 7 Core Marianas™ Microscope equipped with CSU-X1 A1 Spinning Disk Confocal set up and Environmental temperature and CO2 control platform. Images were taken using ORCA Fusion BT sCMOS Camera and analyzed with Slidebook v. 6.0 software. For every time point z-stack was generated (step size 500 nm) and cells were imaged every 10 minutes until completion of the division.

## AUTHOR CONTRIBUTIONS

Experiments were conceived by KFV, AJN, RP, and JMT and were performed by KFV, AJN, FZ, XW, and RP. Funding for the project was secured by KFV, RP, and JMR. The manuscript was written by KFV, RP, and JMT and edited by KFV, AJN, FZ, XW, RP, and JMT.

## RESOURCE AVAILABILITY

### Lead contacts

Further information and requests for resources and reagents should be directed to and will be fulfilled by the lead contacts, Rytis Prekeris and J. Matthew Taliaferro.

### Materials availability

All materials generated in this study are available through the lead contacts upon request.

### Data availability

All high-throughput sequencing data associated with these experiments has been deposited at the Gene Expression Omnibus under accession number GSE342282.

## RESOURCES TABLE

| REAGENT OR RESOURCE | SOURCE | IDENTIFIER |
| --- | --- | --- |
| DMEM | Gibco | CAT# 11965092 |
| Penicillin–streptomycin solution | Gibco | CAT# #15140122 |
| Fetal Reserve Bovine Serum | Atlas Biologicals | CAT# FR-0500A |
| Quick-RNA Miniprep Kit | Zymo Research | CAT# R1054 |
| Quick-RNA Microprep Kit | Zymo Research | CAT# R1050 |
| LunaScript RT Supermix | New England Biolabs | CAT# E3010 |
| PrimeTime® Gene Expression Master Mix | Integrated DNA Technologies | CAT# 1055772 |
| Probe Sets for qPCR | Integrated DNA Technologies |  |
| Oligonucleotide pool | Twist Bioscience |  |
| KOD hot Start Master Mix | Sigma Aldrich | CAT# 71842-3 |
| NucleoMag NGS Clean-up and Size Select | Takara | CAT# 744970.5 |
| pRD-RIPE | Khandelia et al, 2011 (https://pmc.ncbi.nlm.nih.gov/articles/PMC12864922/#CR27) |  |
| BstXI | NEB | CAT# R0113 |
| Zymoclean Gel DNA Recovery Kit | Zymo Research | CAT# D4008 |
| NEB® 10-beta Competent *E. coli* | NEB | CAT# C3019H |
| ZymoPURE Plasmid Maxiprep kit | Zymo Research | CAT# D4203 |
| SuperScript IV Reverse Transcriptase | ThermoFisher | CAT# 18090200 |
| Zymo DNA Clean & Concentrator kit | Zymo Reserach | CAT# D4013 |
| Qubit dsDNA HS Assay Kits | ThermoFisher | CAT# Q32854 |
| Agilent D1000 ScreenTape | Agilent | CAT# 5067-5582 |
| SSC | Fisher Scientific | CAT# 15557044 |
| Formamide | Sigma-Aldrich | CAT# F7503-100ML |
| smFISH hybridization buffer | Biosearch Technologies | CAT #SMF-HB1-10 |
| Fluoromount G | SouthernBiotech | CAT# 0100-01 |
| NET1 antibody | Proteintech | CAT# 28180-1-AP |
| Western blot sample buffer | Invitrogen | CAT# NP008 |
| Spectra Protein Ladder | Thermo Fisher | CAT# 26634 |
| MOPS SDS Running Buffer | ThermoFisher | CAT# NP0001 |
| iBlot2 dry transfer machine | Thermo Fisher | CAT# IB21001 |
| Sapphire imager | Azure Biosystems |  |

---

**Figure S1.**
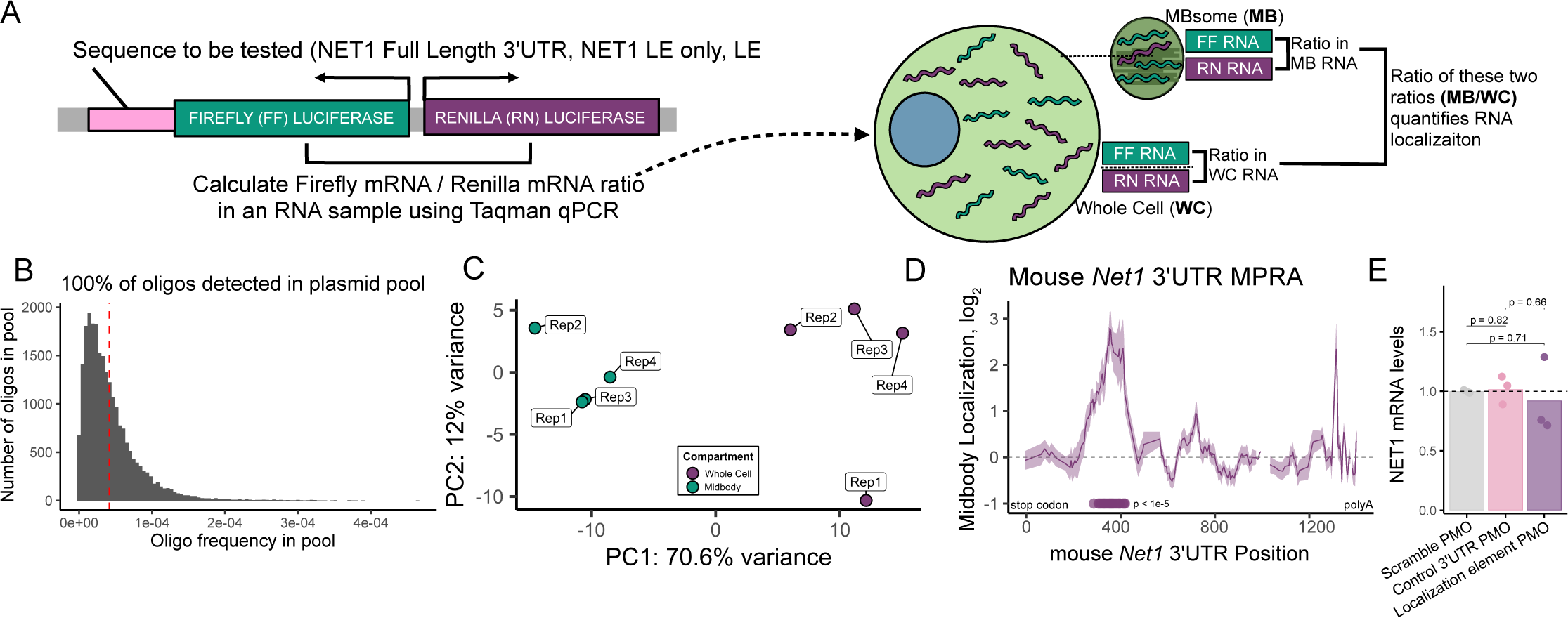
Massively parallel reporter assay and reporter based MB localization assay design. (A) Schematic representation of RT-qPCR-based reporter RNA approach for assaying MB RNA localization. Plasmids expressing Firefly and Renilla luciferases from a bidirectional reporter are integrated into the genome. Sequences to be tested for MB localization activity (*NET1* full length 3′ UTR, LE only and ΔLE 3’UTR) are appended to the 3′ UTR of Firefly luciferase. Using Taqman qPCR, the ratio of Firefly to Renilla luciferase mRNA is measured in MB and whole-cell samples. The ratio of these ratios (MB/WC) quantifies the MB enrichment of the Firefly luciferase transcript, measuring the tested sequence’s effect on MB RNA localization. (B) Distribution of oligonucleotide frequency integrated into the plasmid pool. 100% of oligonucleotides were detected. The red line indicates the expected frequency if all oligonucleotides were equally represented. (C) Principal component analysis representing the difference between 4 whole cell (WC) control and 4 MB RNAseq datasets. Samples separated along PC1 (70.6%) by WC and MB samples. (D) Midbody over whole cell enrichment of oligonucleotides as a function of their location within the mouse *Net1* 3’UTR. The line represents a rolling average of 8 oligonucleotides and the ribbon represents the standard deviation of MB enrichment within the sliding window. Dots below the lines represent 3’UTR positions of significantly MB localized oligos (FDR < 1e-5). The identified localization element is consistent with previously identified mouse Net1 localization element, active in human MBs (Farmer et al. 2023; Arora et al. 2022). (E) Normalized *NET1* mRNA levels following transfection with antisense phosphorodiamidate morpholino oligonucleotides (PMOs) targeting a scrambled control sequence, a control region of the *NET1* 3′UTR, or the *NET1* localization element. Bars represent the mean of individual replicates (dots). P values were calculated using a t-test.

**Figure S2.**
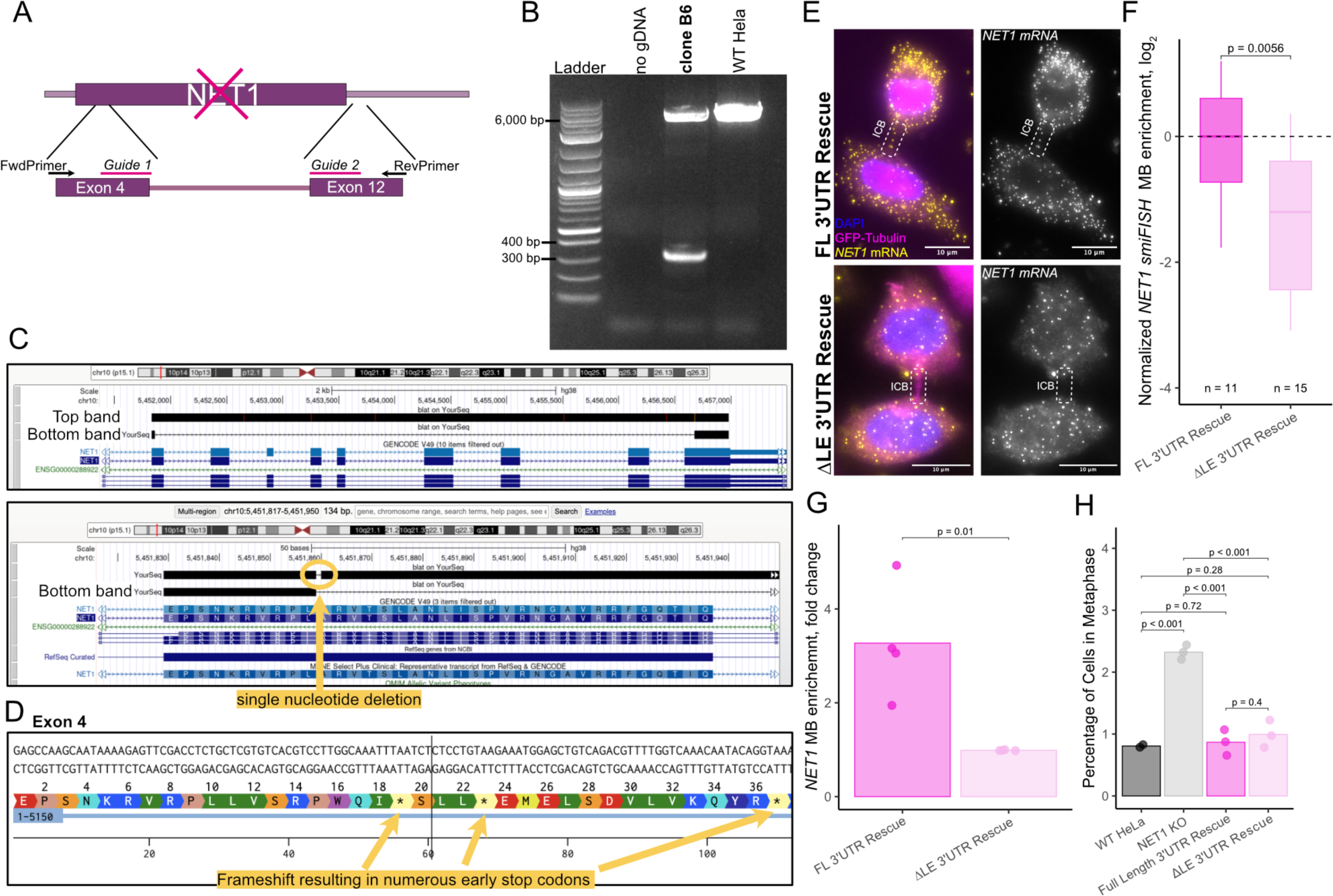
NET1 knockout and rescue validation. (A) Schematic representation of the NET1 genomic locus showing exon 4 (earliest exon shared by all isoforms of NET1) and exon 12 targeted by the CRISPR guide RNAs. The primers shown are used for a genomic DNA PCR to validate the knockout. (B) Genomic DNA PCR of the knockout clone used, compared to the WT gDNA PCR. The bottom band experienced a full dropout of the genomic DNA between the two cute sites, while the top band obtained a single nucleotide deletion in exon 4. (C) Sequencing alignment of the top and bottom PCR bands from NET1 KO clone B6. The top band contains a single nucleotide deletion, while the bottom band contains a full drop out of the genomic sequence between cut sites. (D) The predicted amino acid sequence of the CRISPR edited Exon 4 sequence, with a few of the frameshift-induced premature stop codons indicated. (E) Midbody localization NET1 knockout cells rescued by *NET1* transgenes containing either the full length 3’UTR or the ΔLE 3’UTR (yellow) visualized using smFISH. in cells expressing GFP-tubulin labeling the ICB/MB (pink). ICB is indicated by the dotted box. (F) Quantification of *NET1* mRNA visualized in C. *NET1* puncta in intercellular bridges/MBs and whole cells were quantified for each sample and the ratio between the two locations is reported as normalized midbody enrichment (log_2_). Values were normalized to NET1 KO cells rescued by NET1 FL 3’UTR. P values were calculated using a t-test. (G) Midbody localization of the NET1 rescue constructs quantified using a midbody purification and qPCR (Fig S1A). Values are normalized to the FL 3’UTR rescue. P values were calculated using a t test. (H) Percentage of total cells in metaphase, identified using DAPI and anti-acetylated tubulin antibodies. Each point represents a biological replicate with a minimum of 1000 cells represented.

**Figure S3.**
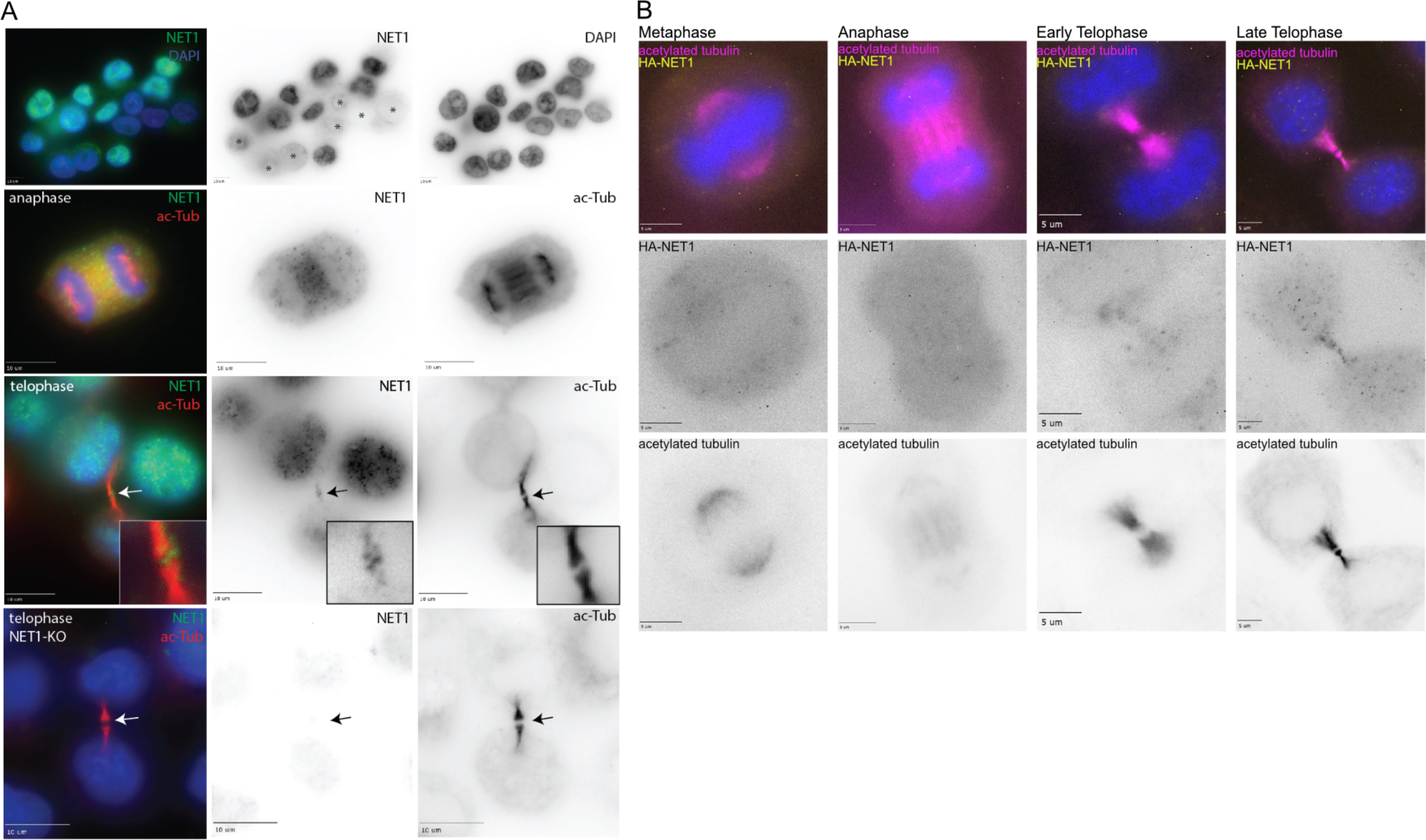
NET1 protein localization. (A) Immunofluorescence staining for endogenous NET1 in wildtype and NET1 knockout HeLa cells. Interphase cells exhibit mostly nuclear localization of NET1. Endogenous NET1 is robustly localized to the ICB/MB (arrow, row 3) in telophase cells. NET1 KO cells exhibit a loss of immunofluorescent signal, shown in a telophase cell completely lacking NET1 at the ICB/MB. (B) HA-NET1 (yellow) localization patterns throughout the stages of mitosis, exhibiting distinct localization to the ICB (visualized using acetylated tubulin stain, magenta) in late telophase.

**Figure S4.**
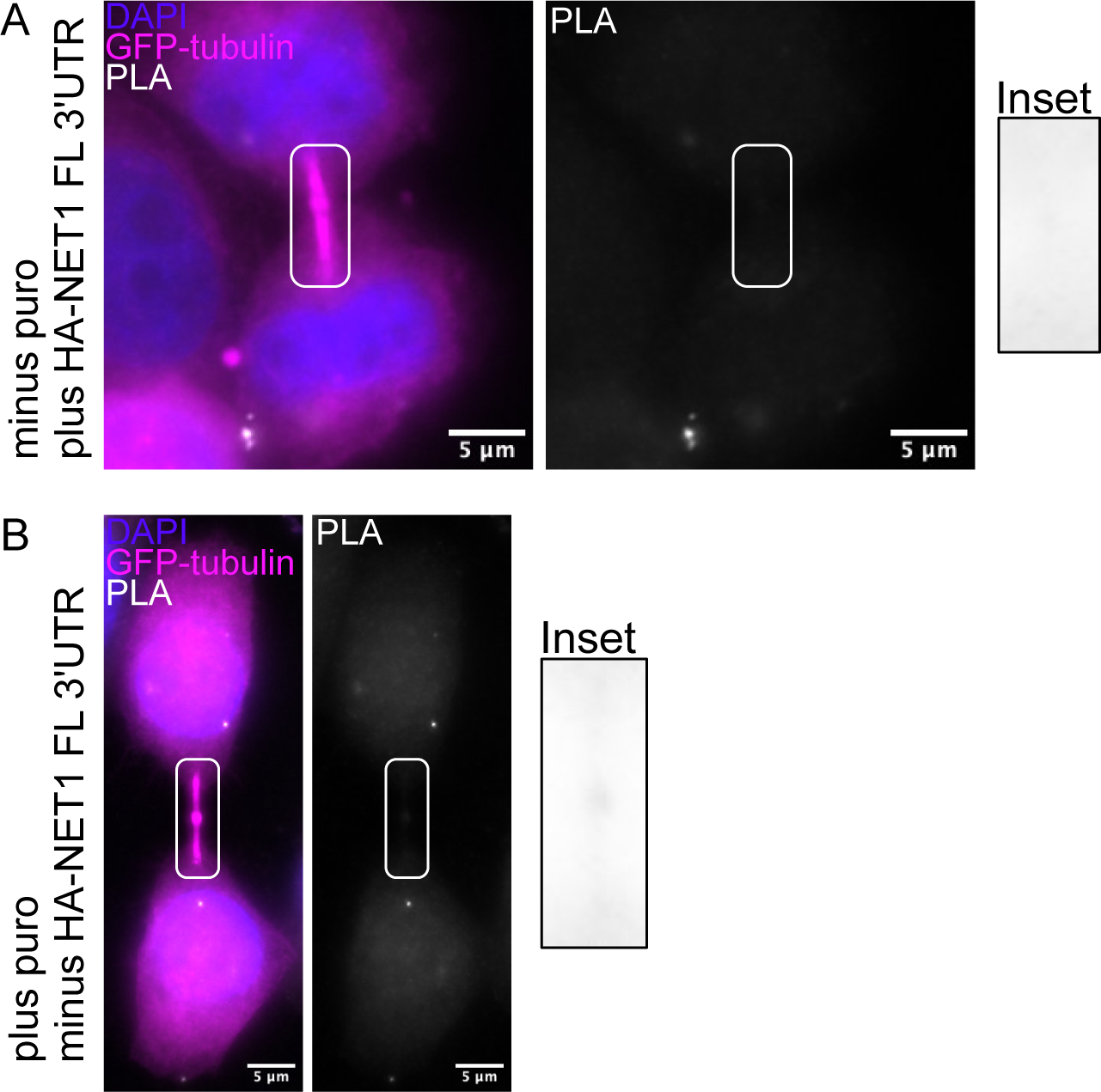
Negative controls for puro-PLA. (A) Representative image for puro-PLA (white) in cells expressing HA-NET1 FL 3’UTR minus puromycin addition. ICB identified by GFP-tubulin (magenta). puro-PLA puncta in ICB shown in the inset. (B) Representative image for puro-PLA (white) in cells not expressing HA-NET1 FL 3’UTR with puromycin addition. ICB identified by GFP-tubulin (magenta). Puro-PLA puncta in ICB shown in the inset.

**Figure S5.**
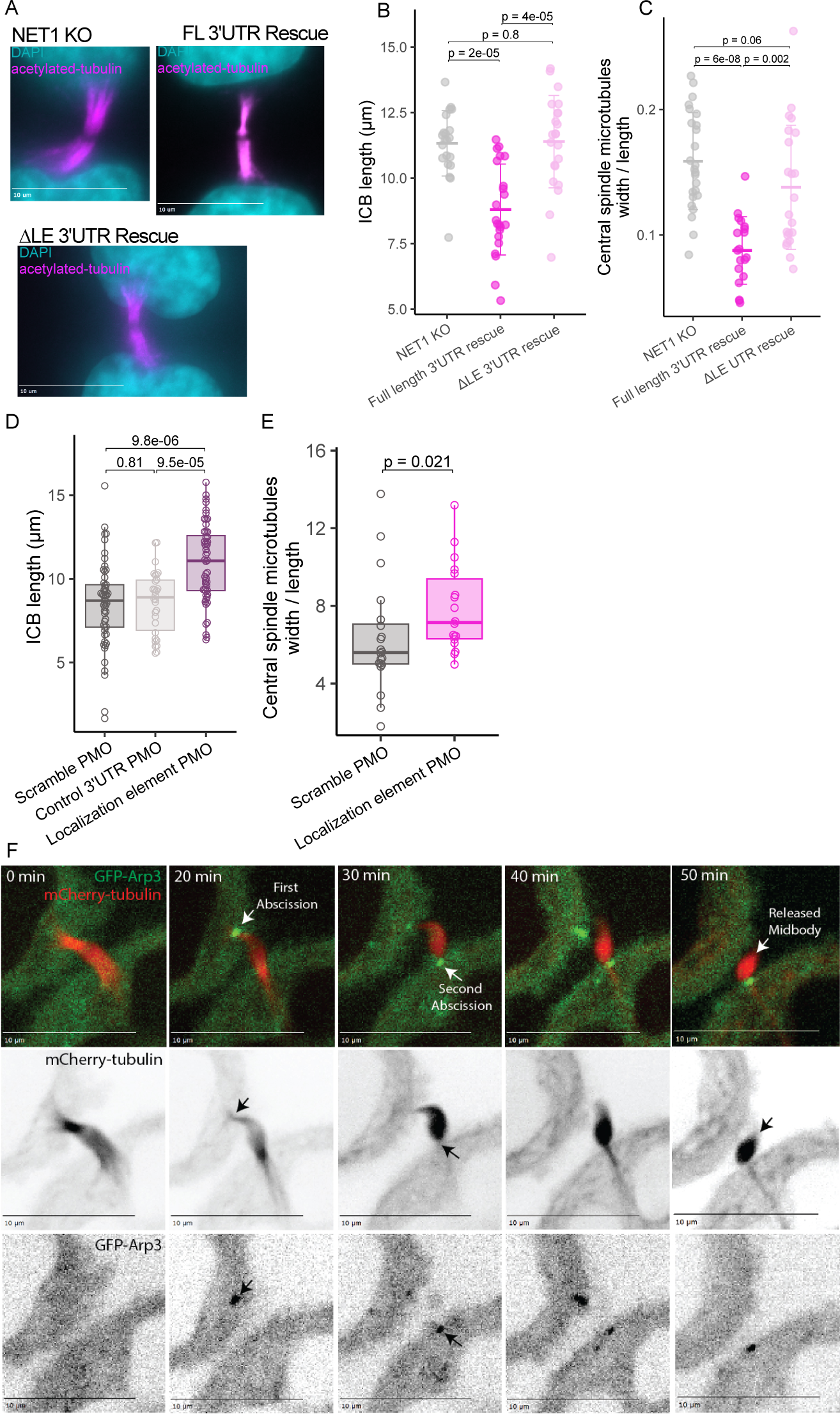
Abscission defects. (A) Representative immunofluorescence images of NET1 knockouts and HA-NET1 rescues in late telophase (ICB labeled by anti- acetylated tubulin antibodies in magenta). NET1 knockout and mis-localization results in elongated and wider bridges. (B) Quantification of ICB length for NET1 KO and HA-NET1 rescues. P values were calculated using a Wilcoxon ranksum test. (C) Quantification of ICB width for NET1 KO and HA-NET1 rescues. Measured as a ratio of central spindle microtubule width over length. P values were calculated using a Wilcoxon ranksum test. (D) Quantification of ICB length for PMO treated cells. P values were calculated using a Wilcoxon ranksum test. (E) Quantification of ICB width for PMO treated cells. Measured as a ratio of central spindle microtubule width over length. P values were calculated using a Wilcoxon ranksum test. (F) Time-lapse live cell imaging of wildtype HeLa cells. Cells expressing GFP-Arp3 (white) and mCherry-tubulin (magenta) were imaged in telophase from right before the first abscission event through the second and final abscission event, showing release of the MBsome after symmetric abscission concludes. Abscission is indicated by the loss of tubulin on one side of the ICB.

**Figure S6.**
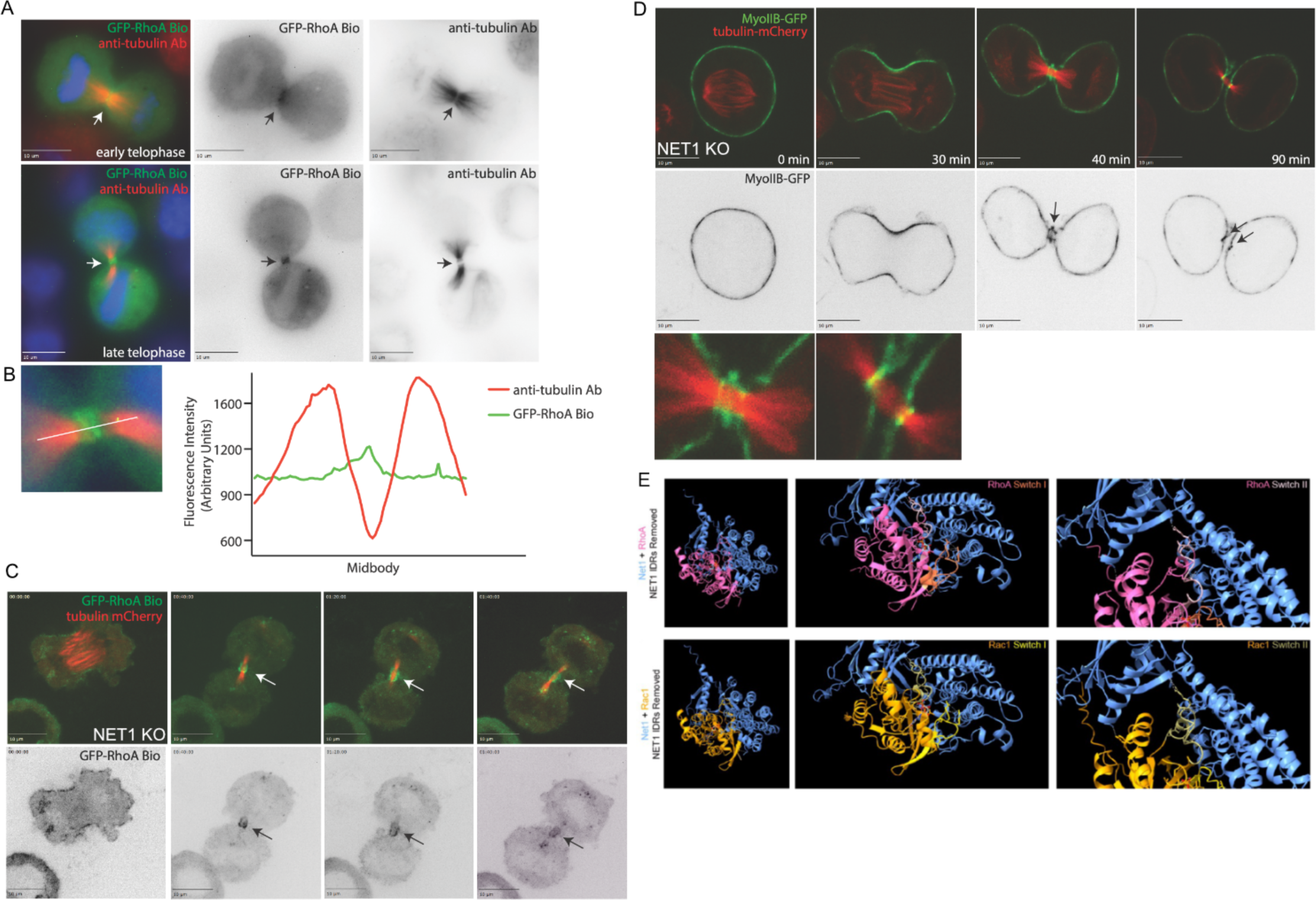
NET1 KO does not affect RhoA activation and Myosin IIB recruitment to the ingressing actomyosin contractile ring. (A) Wildtype HeLa cells expressing GFP-RhoA biosensor (green) stained with anti-acetylated tubulin antibody (red). In early and late telophase RhoA activation occurs throughout the ICB. (B) Line scan analysis of fluorescence intensity for GFP-RhoA biosensor (green) and anti-acetylated tubulin antibody (red) across the ICB. The middle region where loss of tubulin staining occurs indicates the location of the midbody. (C) Time-lapse live cell imaging of NET1 knockout cells. Cells expressing GFP-RhoA biosensor (green) and mCherry-tubulin (red) were imaged through telophase, with no apparent loss of RhoA activation in the ICB or at the MB observed. (D) Time-lapse live cell imaging of NET1 knockout cells. Cells expressing GFP-MyosinIIB, a downstream effector of RhoA, and mCherry-tubulin (red) were imaged from metaphase through telophase, with no apparent loss of MyosinIIB in the ICB. (E) Predicted structural models of NET1 (blue) interaction with Rac1 (yellow, bottom) and RhoA (pink, top). Middle and far right panels show a zoom in of the predicted interfaces at switch I and switch II. RhoA and Rac1 interact similarly, indicating that NET1 might also be a GEF for Rac1 in addition to RhoA.

**Figure S7.**
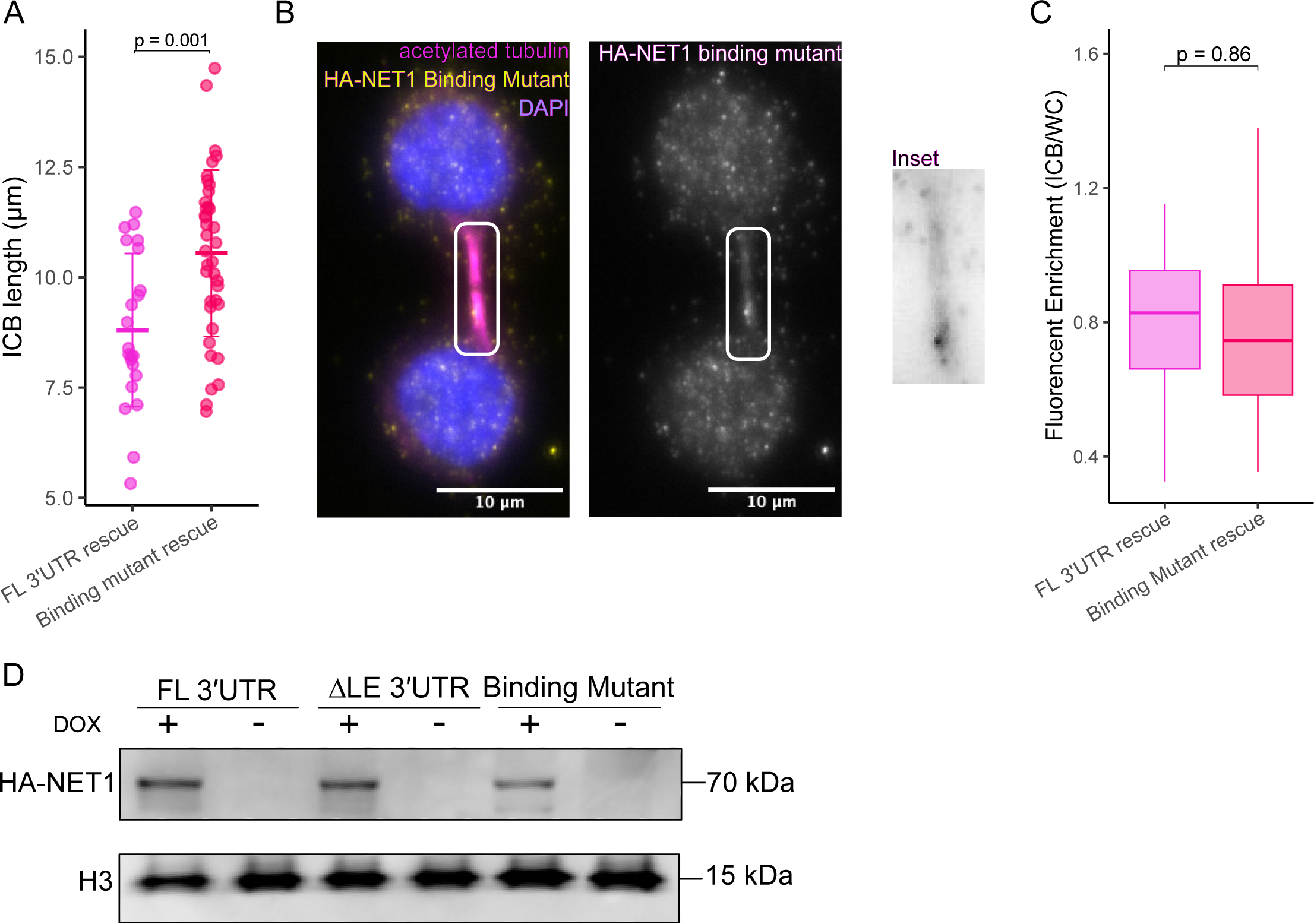
NET1 binding mutant fails to rescue Arp2/3 dependent abscission defect. (A) Quantification of ICB length for HA-NET1 FL 3’UTR and HA-NET1-L321E-FL3’UTR expressing cells. P values were calculated using a Wilcoxon ranksum test. (B) HA-NET1-L321E-FL3’UTR (yellow) localization during late telophase. Cells in late telophase exhibit rounded nuclei (blue), and fully formed ICBs (magenta). ICB indicated by box. (C) Quantification of HA-NET1 enrichment in the ICB as represented by the images in B. P values were calculated using a t-test. (D) Western blot measuring protein expression in HA-NET1-FL 3’UTR, HA-NET1-ΔLE 3’UTR and HA-NET1-L321E-FL3’UTR expressing cell lines with and without doxycycline. Histone 3 (H3) is used as a loading control.

## REFERENCES

1. Addi, Cyril, Adrien Presle, Stéphane Frémont, et al. 2020. “The Flemmingsome Reveals an ESCRT-to-Membrane Coupling via ALIX/syntenin/syndecan-4 Required for Completion of Cytokinesis.” Nature Communications 11 (1): 1941.

2. Advedissian, Tamara, Stéphane Frémont, and Arnaud Echard. 2024. “Cytokinetic Abscission Requires Actin-Dependent Microtubule Severing.” Nature Communications 15 (1): 1949.

3. Alberts, A. S., and R. Treisman. 1998. “Activation of RhoA and SAPK/JNK Signalling Pathways by the RhoA-Specific Exchange Factor mNET1.” The EMBO Journal 17 (14): 4075–4085.

4. Arora, A., R. Castro-Gutierrez, C. Moffatt, et al. 2022. “High-Throughput Identification of RNA Localization Elements in Neuronal Cells.” Nucleic Acids Research 50 (18): 10626–10642.

5. Basant, Angika, and Michael Glotzer. 2018. “Spatiotemporal Regulation of RhoA during Cytokinesis.” Current Biology 28 (9): R570–R580.

6. Bement, William M., Hélène A. Benink, and George von Dassow. 2005. “A Microtubule-Dependent Zone of Active RhoA during Cleavage Plane Specification.” The Journal of Cell Biology 170 (1): 91–101.

7. Carr, Heather S., Christopher A. Morris, Sarita Menon, Eun Hyeon Song, and Jeffrey A. Frost. 2013. “Rac1 Controls the Subcellular Localization of the Rho Guanine Nucleotide Exchange Factor Net1A to Regulate Focal Adhesion Formation and Cell Spreading.” Molecular and Cellular Biology 33 (3): 622–634.

8. Chalamalasetty, Ravindra B., Stefan Hümmer, Erich A. Nigg, and Herman H. W. Silljé. 2006. “Influence of Human Ect2 Depletion and Overexpression on Cleavage Furrow Formation and Abscission.” Journal of Cell Science 119 (Pt 14): 3008–3019.

9. Costa, Guilherme, Joshua J. Bradbury, Nawseen Tarannum, and Shane P. Herbert. 2020. “RAB13 mRNA Compartmentalisation Spatially Orients Tissue Morphogenesis.” The EMBO Journal 39 (21): e106003.

10. Crowell, Elizabeth Faris, Anne-Lise Gaffuri, Barbara Gayraud-Morel, Shahragim Tajbakhsh, and Arnaud Echard. 2014. “Midbody Remnant Engulfment after Cytokinesis Abscission in Mammalian Cells.” Journal of Cell Science, January 1, jcs.154732.

11. Dambournet, Daphné, Mickael Machicoane, Laurent Chesneau, et al. 2011. “Rab35 GTPase and OCRL Phosphatase Remodel Lipids and F-Actin for Successful Cytokinesis.” Nature Cell Biology 13 (8): 981–988.

12. David, Muriel, Dominique Petit, and Jacques Bertoglio. 2012. “Cell Cycle Regulation of Rho Signaling Pathways.” Cell Cycle (Georgetown, Tex.) 11 (16): 3003–3010.

13. D’Avino, Pier Paolo, and Luisa Capalbo. 05/2016. “Regulation of Midbody Formation and Function by Mitotic Kinases.” Seminars in Cell & Developmental Biology 53: 57–63.

14. De Pace, Raffaella, Juan S. Bonifacino, and Saikat Ghosh. 2025. “Puromycin Proximity Ligation Assay (Puro-PLA) to Assess Local Translation in Axons from Human Neurons.” Bio-Protocol 15 (5): e5224.

15. Eden, Sharon, Rajat Rohatgi, Alexandre V. Podtelejnikov, Matthias Mann, and Marc W. Kirschner. 2002. “Mechanism of Regulation of WAVE1-Induced Actin Nucleation by Rac1 and Nck.” Nature 418 (6899): 790–793.

16. Elia, Natalie, Rachid Sougrat, Tighe A. Spurlin, James H. Hurley, and Jennifer Lippincott-Schwartz. 2011. “Dynamics of Endosomal Sorting Complex Required for Transport (ESCRT) Machinery during Cytokinesis and Its Role in Abscission.” Proceedings of the National Academy of Sciences of the United States of America 108 (12): 4846–4851.

17. Engel, Krysta L., Ankita Arora, Raeann Goering, Hei-Yong G. Lo, and J. Matthew Taliaferro. 2020. “Mechanisms and Consequences of Subcellular RNA Localization across Diverse Cell Types.” Traffic 21 (6): 404–418.

18. Farmer, Trey, Katherine F. Vaeth, Ke-Jun Han, Raeann Goering, Matthew J. Taliaferro, and Rytis Prekeris. 2023. “The Role of Midbody-Associated mRNAs in Regulating Abscission.” The Journal of Cell Biology 222 (12): e202306123.

19. Gasparski, Alexander N., Konstadinos Moissoglu, Sandeep Pallikkuth, Sezen Meydan, Nicholas R. Guydosh, and Stavroula Mili. 2023. mRNA Location and Translation Rate Determine Protein Targeting to Dual Destinations. Cell Biology.

20. Goering, Raeann, Ankita Arora, Megan C. Pockalny, and J. Matthew Taliaferro. 2023. “RNA Localization Mechanisms Transcend Cell Morphology.” eLife 12 (March): e80040.

21. Haga, Raquel B., and Anne J. Ridley. 2016. “Rho GTPases: Regulation and Roles in Cancer Cell Biology.” Small GTPases 7 (4): 207–221.

22. Holt, Christine E., and Simon L. Bullock. 2009. “Subcellular mRNA Localization in Animal Cells and Why It Matters.” Science (New York, N.Y.) 326 (5957): 1212–1216.

23. Hu, Chi-Kuo, Margaret Coughlin, and Timothy J. Mitchison. 2012. “Midbody Assembly and Its Regulation during Cytokinesis.” Molecular Biology of the Cell 23 (6): 1024–1034.

24. Jaffe, Aron B., and Alan Hall. 2005. “Rho GTPases: Biochemistry and Biology.” Annual Review of Cell and Developmental Biology 21 (1): 247–269.

25. Khandelia, Piyush, Karen Yap, and Eugene V. Makeyev. 2011. “Streamlined Platform for Short Hairpin RNA Interference and Transgenesis in Cultured Mammalian Cells.” Proceedings of the National Academy of Sciences of the United States of America 108 (31): 12799–12804.

26. Kimura, K., T. Tsuji, Y. Takada, T. Miki, and S. Narumiya. 2000. “Accumulation of GTP-Bound RhoA during Cytokinesis and a Critical Role of ECT2 in This Accumulation.” The Journal of Biological Chemistry 275 (23): 17233–17236.

27. Lawrence, J. B., and R. H. Singer. 1986. “Intracellular Localization of Messenger RNAs for Cytoskeletal Proteins.” Cell 45 (3): 407–415.

28. Love, Michael I., Wolfgang Huber, and Simon Anders. 2014. “Moderated Estimation of Fold Change and Dispersion for RNA-Seq Data with DESeq2.” Genome Biology 15 (12): 550.

29. Macdonald, P. M., and G. Struhl. 1988. “Cis-Acting Sequences Responsible for Anterior Localization of Bicoid mRNA in Drosophila Embryos.” Nature 336 (6199): 595–598.

30. Mahlandt, Eike K., Janine J. G. Arts, Werner J. van der Meer, et al. 2021. “Visualizing Endogenous Rho Activity with an Improved Localization-Based, Genetically Encoded Biosensor.” Journal of Cell Science 134 (17): jcs258823.

31. Martin, Kelsey C., and Anne Ephrussi. 2009. “mRNA Localization: Gene Expression in the Spatial Dimension.” Cell 136 (4): 719–730.

32. Martin, Marcel. 2011. “Cutadapt Removes Adapter Sequences from High-Throughput Sequencing Reads.” EMBnet.journal 17 (1): 10.

33. Mason, Devon E., Thomas D. Madsen, Alexander N. Gasparski, et al. 2025. “Control of Epithelial Tissue Organization by mRNA Localization.” Nature Communications 16 (1): 5216.

34. Menon, Sarita, Wonkyung Oh, Heather S. Carr, and Jeffrey A. Frost. 2013. “Rho GTPase-Independent Regulation of Mitotic Progression by the RhoGEF Net1.” Molecular Biology of the Cell 24 (17): 2655–2667.

35. Mierzwa, Beata, and Daniel W. Gerlich. 2014. “Cytokinetic Abscission: Molecular Mechanisms and Temporal Control.” Developmental Cell 31 (5): 525–538.

36. Neto, Hélia, and Gwyn W. Gould. 2011. “The Regulation of Abscission by Multi-Protein Complexes.” Journal of Cell Science 124 (Pt 19): 3199–3207.

37. Park, Sungjin, Randall Dahn, Elif Kurt, et al. 2023. “The Mammalian Midbody and Midbody Remnant Are Assembly Sites for RNA and Localized Translation.” Developmental Cell 58 (19): 1917–1932.e6.

38. Patel, Smit A., Sungjin Park, Dantong Zhu, et al. 2024. “Extracellular Vesicles, Including Large Translating Vesicles Called Midbody Remnants, Are Released during the Cell Cycle.” Molecular Biology of the Cell 35 (12): ar155.

39. Peterman, E., and R. Prekeris. 2017. “Understanding Post-Mitotic Roles of the Midbody during Cell Differentiation and Polarization.” In Methods in Cell Biology, vol. 137, 137. Elsevier.

40. Peterman, Eric, Paulius Gibieža, Johnathon Schafer, et al. 2019. “The Post-Abscission Midbody Is an Intracellular Signaling Organelle That Regulates Cell Proliferation.” Nature Communications 10 (1): 3181.

41. Piekny, Alisa, Michael Werner, and Michael Glotzer. 2005. “Cytokinesis: Welcome to the Rho Zone.” Trends in Cell Biology 15 (12): 651–658.

42. Qin, Huajun, Heather S. Carr, Xiaochong Wu, Daniella Muallem, Nancy H. Tran, and Jeffrey A. Frost. 2005. “Characterization of the Biochemical and Transforming Properties of the Neuroepithelial Transforming Protein 1.” The Journal of Biological Chemistry 280 (9): 7603–7613.

43. Rossman, Kent L., Channing J. Der, and John Sondek. 2005. “GEF Means Go: Turning on RHO GTPases with Guanine Nucleotide-Exchange Factors.” Nature Reviews. Molecular Cell Biology 6 (2): 167–180.

44. Ryder, Pearl V., and Dorothy A. Lerit. 07/2018. “RNA Localization Regulates Diverse and Dynamic Cellular Processes.” Traffic 19 (7): 496–502.

45. Sachs, Rachel, Yusuke Ogi, and Rytis Prekeris. 2026. “TPGS1 Regulates Central Spindle Microtubule Glutamylation and Remodeling during Telophase and Abscission.” EMBO Reports 27 (8): 1944–1963.

46. Schiel, John A., Glenn C. Simon, Chelsey Zaharris, et al. 2012. “FIP3-Endosome-Dependent Formation of the Secondary Ingression Mediates ESCRT-III Recruitment during Cytokinesis.” Nature Cell Biology 14 (10): 1068–1078.

47. Schmidt, Anja, Joanne Durgan, Ana Magalhaes, and Alan Hall. 2007. “Rho GTPases Regulate PRK2/PKN2 to Control Entry into Mitosis and Exit from Cytokinesis.” The EMBO Journal 26 (6): 1624–1636.

48. Schmidt, Anja, and Alan Hall. 2002. “The Rho Exchange Factor Net1 Is Regulated by Nuclear Sequestration.” The Journal of Biological Chemistry 277 (17): 14581–14588.

49. Skop, Ahna R., Hongbin Liu, John Yates, Barbara J. Meyer, and Rebecca Heald. 2004. “Dissection of the Mammalian Midbody Proteome Reveals Conserved Cytokinesis Mechanisms.” Science 305 (5680): 61–66.

50. Suwakulsiri, Wittaya, Rong Xu, Alin Rai, et al. 2024. “Transcriptomic Analysis and Fusion Gene Identifications of Midbody Remnants Released from Colorectal Cancer Cells Reveals They Are Molecularly Distinct from Exosomes and Microparticles.” Proteomics 24 (11): e2300058.

51. Tatsumoto, T., X. Xie, R. Blumenthal, I. Okamoto, and T. Miki. 1999. “Human ECT2 Is an Exchange Factor for Rho GTPases, Phosphorylated in G2/M Phases, and Involved in Cytokinesis.” The Journal of Cell Biology 147 (5): 921– 928.

52. Tsanov, Nikolay, Aubin Samacoits, Racha Chouaib, et al. 2016. “smiFISH and FISH-Quant – a Flexible Single RNA Detection Approach with Super-Resolution Capability.” Nucleic Acids Research 44 (22): e165–e165.

53. Ulu, Arzu, Wonkyung Oh, Yan Zuo, and Jeffrey A. Frost. 2021. “Cdk1 Phosphorylation Negatively Regulates the Activity of Net1 towards RhoA during Mitosis.” Cellular Signalling 80 (109926): 109926.

54. Wagner, Elizabeth, and Michael Glotzer. 2016. “Local RhoA Activation Induces Cytokinetic Furrows Independent of Spindle Position and Cell Cycle Stage.” The Journal of Cell Biology 213 (6): 641–649.

55. Wang, Jun, Marc Horlacher, Lixin Cheng, and Ole Winther. 2023. “RNA Trafficking and Subcellular Localization—a Review of Mechanisms, Experimental and Predictive Methodologies.” Briefings in Bioinformatics 24 (5): bbad249.

56. Wang, Shiwei, Xuan Wu, Mengmeng Zhang, et al. 2024. “NET1 Is a Critical Regulator of Spindle Assembly and Actin Dynamics in Mouse Oocytes.” Reproductive Biology and Endocrinology: RB&E 22 (1): 5.

57. Yüce, Ozlem, Alisa Piekny, and Michael Glotzer. 2005. “An ECT2-Centralspindlin Complex Regulates the Localization and Function of RhoA.” The Journal of Cell Biology 170 (4): 571–582.

